# Tousled-like kinase activity is required for transcriptional silencing and suppression of innate immune signaling

**DOI:** 10.1101/621409

**Authors:** Sandra Segura-Bayona, Marina Villamor-Payà, Camille Stephan-Otto Attolini, Travis H. Stracker

## Abstract

The Tousled like kinases 1 and 2 (TLK1/2) control histone deposition through the ASF1 histone chaperones and are regulated by the DNA damage response. Depletion of TLK activity caused replication stress, increased chromosomal instability and cell arrest or death. Here, we show that stalled forks in TLK depleted cells are processed by BLM, SAMHD1 and the MRE11 nuclease to generate ssDNA and activate checkpoint signaling. TLK depletion also impaired heterochromatin maintenance, inducing features of alternative lengthening of telomeres and increasing spurious expression of other repetitive elements, associated with impaired deposition of the histone variant H3.3. TLK depletion culminated in a BLM-dependent, STING-mediated innate immune response. In many human cancers, TLK1/2 expression correlated with signatures of chromosomal instability and anti-correlated with STING and innate and adaptive immune response signatures. Together, our results show that TLK activity protects replication forks from active processing, contributes to chromatin silencing and suppresses innate immune responses, suggesting that TLK amplification may protect chromosomally unstable cancers from immune detection.

## Introduction

At early stages in carcinogenesis, oncogene-driven unscheduled DNA replication causes replication stress, a state characterized by slowed or stalled replication forks, increased levels of single-stranded DNA (ssDNA) and activation of the DNA damage response (DDR) (Bartkova et al., 2005; Gorgoulis et al., 2005). DDR signaling acts as an inducible barrier to cancer progression, setting up a selective pressure for tumors to bypass the DDR. Unscheduled S-phase entry may generate an environment where nucleotides, replication factors, histones or other resources are limiting (Bester et al., 2011; Halazonetis et al., 2008; Lee et al., 2018b). In addition, oncogene overexpression can induce the firing of replication origins within highly transcribed genes, leading to replication-transcription conflicts that can provoke chromosomal rearrangements (Kotsantis et al., 2018; Macheret and Halazonetis, 2018). Tumor cells with high basal levels of chromosomal instability (CIN) and replication stress depend on the activity of the ATR and CHK1 kinases, and likely numerous other factors, to maintain CIN and replication stress at sub-toxic levels (Murga et al., 2011; Toledo et al., 2011).

The dynamic maintenance of chromatin structure is required for maintaining both genome and epigenome stability and regulating cell fate (Alabert and Groth, 2012; Yadav et al., 2018). ATR/CHK1-mediated DDR signaling is linked to the maintenance of chromatin structure through the regulation of Tousled like kinase activity (TLK) (Groth et al., 2003; Krause et al., 2003). The Tousled like kinases, TLK1 and TLK2, are conserved Ser-Thr kinases that are critical for the regulation of ASF1a and ASF1b, histone H3/H4 chaperones with key roles in histone deposition during DNA replication, DNA repair and transcription (Carrera et al., 2003; Han et al., 2003; Pilyugin et al., 2009; Roe et al., 1993; Sillje and Nigg, 2001; Sillje et al., 1999). Depletion of both ASF1a and ASF1b results in the arrest of replication forks in the absence of ssDNA generation or replication stress signaling, even in cells treated with the ribonucleotide reductase inhibitor hydroxyurea (HU) or DNA polymerase alpha-primase inhibitor aphidicolin (APH), that normally induce high levels of replication stress (Groth et al., 2007). Depletion of total ASF1 has also been shown to induce features of Alternative lengthening of telomeres (ALT), a telomerase-independent, recombination based telomere maintenance mechanism used in 10-15% of tumors (O’Sullivan et al., 2014; Sobinoff and Pickett, 2017). ALT+ tumors are frequently characterized by inactivation of the ATRX-DAXX complex, which is involved in histone H3.3 deposition and the maintenance of pericentromeric and telomeric heterochromatin. ASF1 has been shown to facilitate H3.3 deposition through the HIRA or CAF1 chaperones, which may be required to compensate for the lack of ATRX-DAXX in ALT+ tumors (Clement et al., 2018; Lovejoy et al., 2012).

In cancer cells, defective chromatin assembly caused by depletion of ASF1B limits cellular proliferation and high levels of ASF1B correlate with poor patient outcome in breast cancer (Corpet et al., 2011). Similarly, Tousled like kinases (TLKs) are critical for nucleosome assembly during DNA synthesis and for replication fork stability, but in contrast to ASF1 depletion that arrests forks without provoking ssDNA accumulation, their loss leads to acute replication stress, characterized by increased levels of ssDNA and DNA double-strand breaks (DSBs) (Groth et al., 2007; Lee et al., 2018b; Mejlvang et al., 2014). Both *TLK1* and *TLK2* are maintained in most cancers and often amplified. In some cases their increased expression correlates with poor prognosis, suggesting that they may be potential cancer targets (Kim et al., 2016; Lee et al., 2018b).

DNA damage inducing chemotherapy elicits an inflammatory response involving the induction and secretion of cytokines (Li and Chen, 2018; Rodier et al., 2009). This response relies on the detection of cytoplasmic DNA by the cGAS protein that activates the Stimulator of Interferon Genes (STING, encoded by *TMEM173*) protein. Several nuclease activities, including TREX1, SAMHD1 and MRE11 have been implicated in the suppression of the interferon (IFN) response (Pasero and Vindigni, 2017). SAMHD1 in particular localizes to nuclear replication forks where it recruits the MRE11 nuclease to regulate replication fork progression and prevent IFN activation in response to replication stress (Coquel et al., 2018). In addition to the so far poorly defined substrates generated at replication forks, both micronuclei, which can result from replication stress, and extrachromosomal single-stranded C-rich (CCCTAA) telomeric circles (C-circles) that are produced in ALT+ cancer cells, are potent inducers of cGAS-STING signaling (Chen et al., 2017; Harding et al., 2017; Mackenzie et al., 2017).

As our previous results indicated that the depletion of TLKs caused replication stress, chromosomal instability and cell arrest or death (Lee et al., 2018b), we sought to better understand the origin of these phenotypes and the mechanisms leading to toxicity in TLK depleted cells. Here, we show that replication stress signaling following the depletion of TLKs is dependent on BLM, SAMHD1 and MRE11 nuclease activity and often occurs in the proximity of telomeres, natural sites of replication stress (Sfeir et al., 2009). In ALT+ U-2-OS cells, most replication stress signaling co-localizes with telomeric markers and similar to ASF1 depletion, TLK depletion in ALT-HeLa LT cells induces hallmarks of ALT, including increased telomeric extrachromosomal C-circles. TLK depletion leads to increased chromatin accessibility at heterochromatin, reduced H3.3 deposition and a robust induction of STING-mediated innate immune responses, regardless of treatment with replication stress inducing agents or SAMHD1-dependent DDR signaling. Finally, in human tumors, *TLK1* and *TLK2* expression correlated with CIN and anti-correlated with innate and adaptive immune responses in many tumor types. Our results indicate that TLK activity is critical for replication fork protection and heterochromatin maintenance, suggesting that TLK amplification in cancer may prevent the innate immune response induced by CIN and ALT, thus limiting immune recognition of tumors.

## Results

### Replication stress signaling following TLK depletion is helicase/nuclease dependent

ASF1 is required for ssDNA generation following replication stress induced by treatment with hydroxyurea (HU) or Aphidicolin (APH) (inhibitors of ribonucleotide reductase and polymerase alpha, respectively) and was proposed to facilitate the generation of ssDNA by facilitating duplex unwinding by the replicative helicase due to uncoupling from the stalled polymerase (Figure S1A) (Groth et al., 2007). To determine if this required TLK activity, we analyzed ssDNA generation following HU or APH treatment of TLK depleted cells by measuring chromatin-bound RPA levels. In contrast to what has been observed for ASF1 depletion, HU treatment led to extensive replication stress signaling in TLK1/2 depleted cells, albeit chromatin-bound RPA levels were slightly reduced compared to siRNA controls, in line with the reduced fork speed in TLK depleted cells (Figure 1A and 1B) (Lee et al., 2018b). Release of cells into fresh media caused higher levels of ssDNA accumulation in TLK1/2 depleted cells and upon continuous treatment with low dose HU or APH, more ssDNA and DSB signaling was observed upon TLK loss (Figure 1A and S1B). This also rendered TLK1/2 depleted cells more sensitive to both HU and APH treatment (Figure 1C) consistent with their higher levels of replication stress signaling and DNA damage (Figure 1A, 1B and S1B).

**Figure 1.**
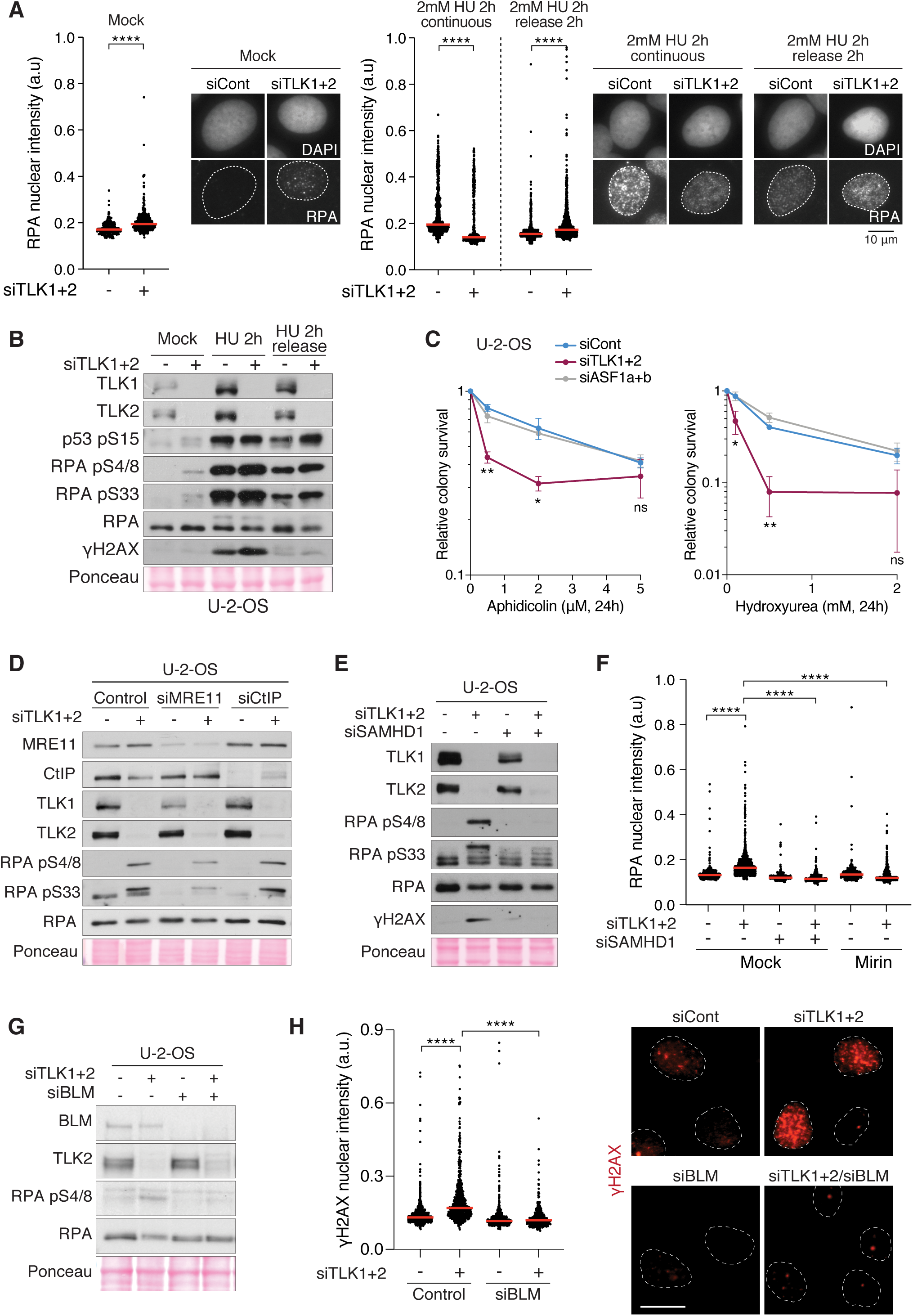
Replication stress signalling following TLK depletion is dependent on nuclease degradation of regressed forks. **(A)** High throughput microscopy (HTM) quantification of the nuclear intensity of chromatin-bound RPA in U-2-OS cells 48 hours after siRNA transfection mock treated or upon the addition of 2 mM Hydroxyurea for 2 hours and harvested without a release or with a release in fresh media for 2 hours. Red bars indicate the median. **(B)** Western blot showing depletion of TLKs in U-2-OS cells treated as in (A). Ponceau staining is shown as a loading control. **(C)** Sensitivity of TLK and ASF1 depleted cells to Hydroxyurea and Aphidicolin measured by colony formation assay. Means and range of two biological replicates performed in technical duplicate are shown. **(D)** Western blot showing single or double depletion of TLKs and MRE11/CtIP in U-2-OS cells harvested 48 hours after siRNA transfection. Ponceau staining is shown as a loading control. **(E)** Western blot showing single or double depletion of TLKs and SAMHD1 in U-2-OS cells harvested 48 hours after siRNA transfection. Ponceau staining is shown as a loading control. **(F)** HTM quantification of the nuclear intensity of chromatin-bound RPA in U-2-OS cells 48 hours after siRNA transfection, mock treated or treated with 50 µM Mirin for 5 hours prior to harvesting. Red bars indicate the median. **(G)** Western blot showing single or double depletion of TLKs and BLM in U-2-OS cells harvested 48 hours after siRNA transfection. Ponceau staining is shown as a loading control. **(H)** HTM quantification of the nuclear intensity of chromatin-bound γH2AX in U-2-OS cells 48 hours after siRNA transfection. Red bars indicate the median. Representative images are shown in the right panel. ****P < 0.0001, ***P < 0.001, **P < 0.01, *P < 0.05, unpaired t test with Welch’s correction (Figure 1A, F and H), unpaired t test (Figure 1C)

In addition to unwinding, ssDNA can be generated by helicases or nuclease-mediated resection of the fork, particularly when it is regressed (Figure S1A) (Quinet et al., 2017). To determine if stalled forks in TLK depleted cells were actively targeted by resection enzymes, we co-depleted or inhibited several candidate proteins, including MRE11, CTIP, SAMHD1 and the BLM helicase. MRE11 mediates DNA end resection at stalled replication forks, in conjunction with either the CtIP or SAMHD1 proteins that facilitate its recruitment and activity, and BLM has been shown to facilitate the resection activities of other nucleases including EXO1 (Coquel et al., 2018; Gravel et al., 2008; Nimonkar et al., 2011; Sartori et al., 2007). Depletion of MRE11 or SAMHD1 with siRNA or inhibition of MRE11 with Mirin strongly reduced replication stress signaling (p-RPA and γH2AX) and the chromatin bound RPA signal detected in TLK depleted cells (Figure 1D, 1E, 1F and S1C) (Dupre et al., 2008). In contrast, depletion of CtIP did not influence replication stress signaling in TLK depleted cells, (Figure 1D). Similarly to MRE11 or SAMHD1 depletion, siRNA depletion of BLM abrogated ssDNA and DNA damage accumulation (Figure 1G and 1H). These data indicated that TLK1/2 activity was not required for ASF1-dependent generation of ssDNA following HU or APH treatment and showed that BLM, MRE11 and SAMHD1 are required to generate ssDNA following TLK depletion.

### Loss of TLK activity compromises heterochromatin maintenance

Given that TLK depletion reduced *de novo* histone deposition and provoked replication stress and sensitivity to ATR-CHK1 inhibitors, HU and APH (Figure 1) (Lee et al., 2018b), we sought to determine if particular genomic regions, such as fragile sites or telomeres (Casper et al., 2002; Flynn et al., 2015), may be hypersensitive to TLK1/2 depletion. In order to map the genome-wide effects of TLK depletion on chromatin accessibility, we used the Assay for Transposase-Accessible Chromatin (ATAC) (Buenrostro et al., 2013) followed by deep-sequencing (ATAC-seq) or by qPCR (ATAC-qPCR). We first evaluated chromatin accessibility in U-2-OS, following depletion of TLK1, TLK2 or both TLK1/2. We considered consensus peaks as those that appeared in at least one of our four conditions, so we could identify relative chromatin accessibility changes (Figure 2A). From consensus peaks covering the whole annotated genome, 10% of the genome became differentially accessible upon total TLK depletion (Figure 2B). These significant changes involved regions depleted of accessibility peaks (negative FC), which were mainly promoter-TSS regions, and regions enriched with accessibility peaks (positive FC), which were mainly intronic and intergenic regions (Figure 2B, 2C, S2A and S2B). Single depletion of TLK1 or TLK2 caused a similar trend, but differences were always more evident following the depletion of both TLK1/2 (Figure S2B).

**Figure 2.**
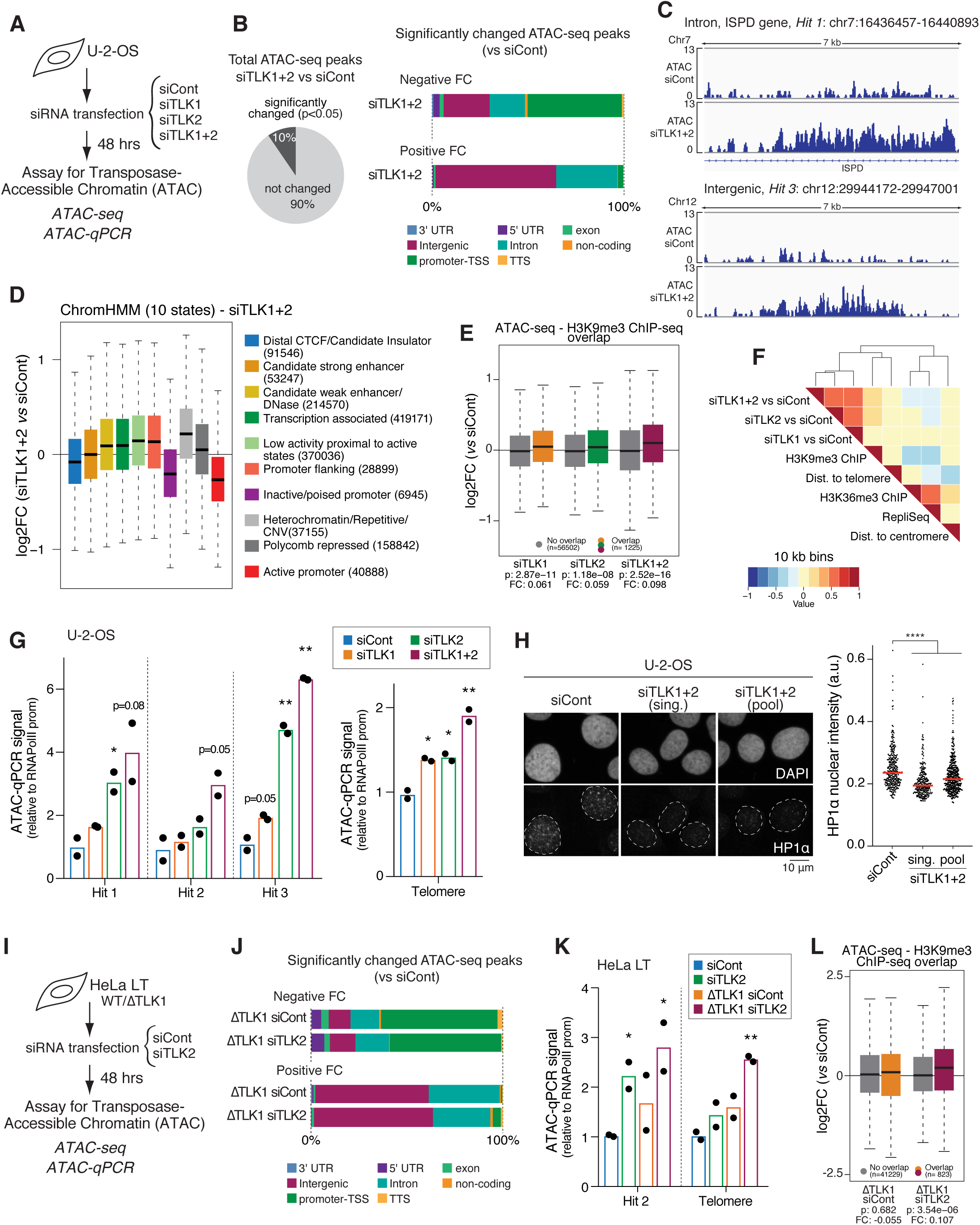
Loss of TLK activity compromises heterochromatin maintenance. **(A)** Schematic depicting the experimental design of ATAC experiment in U-2-OS cells. **(B)** Summary of changes on chromatin accessibility in U-2-OS upon depletion of TLK1/2 are statistically significant in 10% of the genome (n=2) (left panel). Genome annotation of peaks with statistically significant negative or positive fold change upon TLK depletion (right panel). **(C)** Representative IGV tracks of ATAC-seq reads of an intronic region (top panel) and intergenic region (bottom panel) that become more accessible upon TLK loss. **(D)** Boxplots of ATAC-seq FC (siTLK1+2 relative to siCont) through different ChromHMM chromatin states. **(E)** Boxplots depicting ATAC-seq FC relative to siCont computed in regions that do or do not overlap with H3K9me3 ChIP-seq (GSM788078). **(F)** Heatmap that shows the correlation relationship grouped using hierarchical clustering between ATAC-seq FC (relative to siCont) and large-scale chromatin features at 10 kb resolution. **(G)** ATAC-qPCR was used to assess open chromatin at selected genomic regions in U-2-OS cells. Data of qPCR amplification were normalized to unchanging genomic region (RNAPol II promoter region) and represented as the fold-change relative to the signal obtained in siCont conditions, which were set to 1 (n=2). **(H)** Representative IF images of HP1α staining in U-2-OS cells (left panel). HTM quantification of chromatin-bound HP1α levels (right panel), data are from one biological replicate with n>300 nuclei analysed, and representative of four biological replicates. Median is shown in red. **(I)** Schematic depicting the experimental design of ATAC experiment in HeLa LT cells. **(J)** Genome annotation of peaks with statistically significant negative or positive fold change upon TLK depletion (n=2). **(K)** ATAC-qPCR at selected genomic regions in HeLa LT cells as in (G) (n=2). **(L)** Boxplots depicting ATAC-seq FC relative to siCont computed in regions that do or do not overlap with H3K9me3 ChIP-seq (GSM788078). ****P < 0.0001, ***P < 0.001, **P < 0.01, *P < 0.05, unpaired t test with Welch’s correction (Figure 2H), unpaired t test (Figure 2G, K)

We focused further analysis on regions that became more accessible, as this could reflect defective chromatin assembly (Figure 2C). To evaluate how accessibility changes correlated with particular genomic and epigenomic features, we assessed the statistical overrepresentation of ATAC-seq results using diverse published datasets. We correlated our fold changes with chromatin states classified by the ChromHMM tool (Ernst and Kellis, 2012) and observed that heterochromatin (in particular constitutive heterochromatin defined as repetitive/CNV) had the most prominent increase in accessibility upon TLK loss (Figure 2D). Consistent with this, we found that ATAC peaks overlapping with the H3K9me3 histone modification, a marker of heterochromatin (ChIP-seq dataset GSM788078), were enriched in TLK depleted samples (Figure 2E). At a medium resolution of 10 kb, increased accessibility in TLK depleted cells was positively correlated with the heterochromatic mark H3K9me3 (Figure 2E and 2F) and negatively correlated with the euchromatin mark H3K36me3 and early replication timing (Figure 2F and S2C) At this resolution, we did not observe any correlation with chromosomal locations including distance to telomeres or centromeres (Figure 2F). Validation by ATAC-qPCR of some of the higher FC heterochromatic peaks (hit 1 to hit 3), as well as telomeres, confirmed their increased accessibility upon TLK loss (Figure 2G). To determine if TLK depletion had a general impact on heterochromatin maintenance, we examined the chromatin-bound pool of HP1α, a marker of pericentric heterochromatin, and observed a strong reduction in HP1α signal in quantitative immunofluorescence following TLK depletion (Figure 2H).

Given that U-2-OS cells are ALT+ and lack the H3.3 chaperone ATRX, we considered that they could be more dependent upon TLK-ASF1 for H3.3 deposition that is required for heterochromatin maintenance, thus explaining the pronounced reduction of HP1α (Figure 2H). We therefore evaluated chromatin accessibility in HeLa long telomere (LT) cells, a clone of the HeLa 1.2.11 cell line that was derived with long telomeres of about 20 kb (Figure 2I) (O’Sullivan et al., 2014) and found that while the significance of the changes was minor-only 2-5% of consensus peaks displayed statistical significance-the genomic annotation pattern of peaks that were enriched or depleted upon TLK loss was very similar to that of U-2-OS (Figure 2J and S2D). We further validated hit regions and telomeres by ATAC-qPCR in HeLa LT cells (Figure 2K and S2E) and noted that ATAC peaks overlapping with H3K9me3 mark were enriched in ΔTLK1 siTLK2 samples (Figure 2L). Overall, these results supported that constitutive heterochromatin, which is restricted mainly to gene-poor regions, pericentromeres and telomeres, comprised the chromatin regions most dependent on TLK activity for maintenance. Moreover, our findings imply that ALT+ U-2-OS cells, lacking the functional H3.3 chaperone ATRX-DAXX, may be particularly dependent on TLK activity for heterochromatin maintenance, as the ASF1-mediated delivery of H3.3 to the HIRA chaperone is the only known route for H3.3 deposition in these cells.

### TLK activity promotes H3.3 deposition and suppresses telomeric recombination

To further address this possibility, we investigated the effects of TLK depletion on H3.3 deposition in more detail. Analysis of H3.3 ChIP-seq upon TLK depletion in U-2-OS cells resulted in an average decrease of H3.3 occupancy in all genomic regions examined, including TSS regions (Figure 3A, S3A and S3B). Histone H3.3 is required for the establishment of H3K9me3 and formation of heterochromatin and its absence generates DNA damage and increased telomeric sister chromatid exchange (tSCE) in mouse ES cells (Udugama et al., 2015). We observed that in the absence of TLKs, H3.3 and H3K9me3 occupancy was decreased at telomeres (Figure 3B) and expression of telomeric repeat-containing RNA (TERRA) transcripts was elevated (Figure 3C), suggesting it may compromise heterochromatin and promote hyper-recombination at these sites.

**Figure 3.**
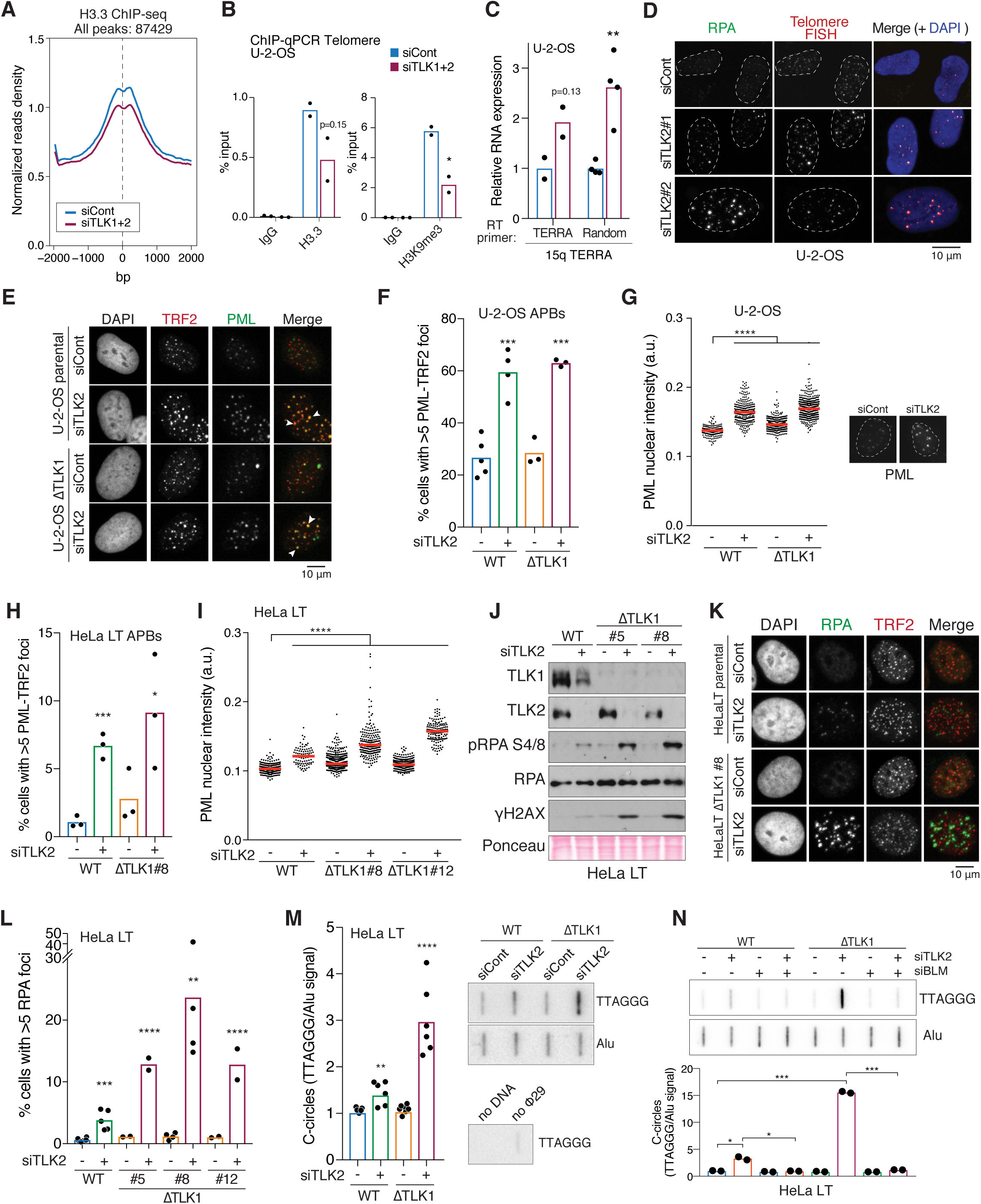
TLK activity promotes H3.3 deposition and suppresses telomeric recombination. **(A)** Normalized H3.3 ChIP-seq read density at all centred H3.3 peaks with +/− 2 kb performed in U-2-OS cells (n=2). **(B)** ChIP-qPCR was used to assess occupancy of H3.3 and H3K9me3 at telomeres in U-2-OS cells. Data of qPCR amplification were normalized to input (n=2). **(C)** Expression of 15q TERRA by RT-qPCR in U-2-OS cells. Data were normalized to unchanging expression gene levels (B-actin) and the signal obtained in siCont conditions was set to 1 (n=2 for RT primer “TERRA”, n=4 for RT primer “Random”). **(D)** IF-FISH staining of chromatin bound RPA and telomeres in U-2-OS cells. **(E)** Representative IF of APBs in U-2-OS cells. White arrowheads indicate colocalization of TRF2-PML foci. **(F)** Quantification of APBs in U-2-OS cells scored as cells with more than 5 TRF2-PML colocalizing foci, with more than 100 cells scored per individual experiment (n=5 for siCont, n=4 for siTLK2, n=3 for ΔTLK1 siCont/siTLK2). **(G)** HTM quantification of nuclear intensity of chromatin-bound PML bodies in U-2-OS cells. Data are from one biological replicate with n>180 nuclei analysed, and representative of four biological replicates. Representative images are shown. Median is shown in red. **(H)** Quantification of APBs in HeLa LT cells, scored as in (F), with more than 100 cells scored per individual experiment (n=3). **(I)** HTM quantification of chromatin-bound PML nuclear intensity in HeLa LT cells, data are from one biological replicate with n>100 nuclei analysed, and representative of three biological replicates. Median is shown in red. **(J)** Western blot of replication stress and DNA damage signalling markers upon TLK2 depletion in HeLa LT cells, parental (WT) and TLK1 CRISPR knockout clones (ΔTLK1). **(K)** Representative IF of RPA-TRF2 staining in HeLa LT cells. **(L)** Quantification of cells with more than 5 RPA discrete foci, with more than 200 cells evaluated per individual experiment (n=6 siCont, n=5 siTLK2, n=2 ΔTLK1#5 and ΔTLK1#12, n=4 ΔTLK1#8). **(M)** Quantification of telomeric C-circles in HeLa LT cells WT or ΔTLK1 72h after being treated with the corresponding siRNAs. Telomeric signal (TTAGGG) was normalized by Alu signal (n=7 siCont, n=6 siTLK2). Representative C-circle assay dot blot is shown in the right panel. **(N)** Quantification of telomeric C-circles in HeLa LT cells as in (N). Telomeric signal (TTAGGG) was normalized by Alu signal (n=2). Representative C-circle assay dot blot is shown in the top panel. ****P < 0.0001, ***P < 0.001, **P < 0.01, *P < 0.05, unpaired t test with Welch’s correction (Figure 3G, I), unpaired t test (Figure 3B-C, F, H, L, N), one-sample t test (Figure 3M)

To address this further, we cytologically analyzed the relationship between replication stress signaling and telomeres in U-2-OS cells. Nearly all of the RPA accumulated in discrete foci that co-stained with the shelterin subunit TRF2 or telomeric FISH signal upon depletion of TLK activity (Figure 3D, S3C and S3D). The increased accumulation of RPA at telomeres suggested the exposure of telomeric ssDNA, indicating the presence of potentially recombinogenic products. TLK2 or total TLK depletion in U-2-OS cells caused increased numbers of ALT-associated PML bodies (APBs) addressed by co-staining with TRF2 and PML (Figure 3E and 3F). In addition, we noticed a remarkable increase in the fluorescent intensity of these PML bodies, providing evidence for upregulation of PML and potential telomere clustering (Figure 3G) (Draskovic et al., 2009).

As ASF1 depletion was shown to induce the features of ALT in ALT-cell lines (O’Sullivan et al., 2014), we next examined these effects in HeLa LT cells where telomere elongation occurs via telomerase. Total TLK depletion, achieved by depleting TLK2 in TLK1 knockout clones (ΔTLK1), increased the presence of ALT-associated PML bodies and resulted in a prominent increase in the fluorescent intensity of PML bodies (Figure 3H and 3I). Total TLK loss caused replicative stress and DNA damage in these cells, visualized in Western blot by phosphorylation of RPA at S4/8 and γH2AX (Figure 3J) and in immunofluorescence by accumulation of chromatin-bound RPA that often co-localized with telomeres, albeit less dramatically than in U-2-OS (Figure 3K, 3L and S3E). In order to assess recombination at ALT-telomeres, we quantified the presence of C-circles that are generated as a byproduct of ALT (Henson et al., 2009; Henson et al., 2017). Following the loss of both TLKs, production of C-circles was significantly enhanced by about 3-fold relative to control conditions (Figure 3M). Moreover, C-circle production was dependent on BLM, as its depletion abrogated C-circle production (Figure 3N and S3F). Together, these results indicated that in the absence of a functional ATRX-DAXX pathway, cells may rely on the TLK-ASF1 pathway for histone H3.3 maintenance at telomeres and other heterochromatic sites. This is substantiated by the fact that similar, but less dramatic phenotypes, were observed in HeLa LT cells that express ATRX and telomerase.

### TLK depletion compromises silencing and activates innate immune signaling

In addition to the effects observed at telomeres, disruption of constitutive heterochromatin formation, which is largely transcriptionally repressed and comprises mostly repetitive DNA, could affect the silencing of other well-established heterochromatic sites. To address this, we performed RNA-seq with rRNA depletion in U-2-OS cells, so that apart from mRNA analysis we could study other RNA species devoid of a poly(A) tail. We first analyzed differential RNA expression of samples by aligning to GENCODE annotated genetic variants (Harrow et al., 2012), which include protein-coding loci with alternatively spliced variants, non-coding loci and pseudogenes. From the categories annotated in GENCODE, depletion of TLKs only significantly affected the expression of antisense RNA and long intergenic noncoding RNA (lincRNA). Both antisense and lincRNA transcripts were upregulated upon TLK loss (Figure 4A) and the double depletion of both TLK1 and TLK2 caused an additive phenotype, consistent with both TLKs independently contributing to the suppression of noncoding RNA transcription. By aligning our RNA-seq data to the annotated database for repetitive elements Repbase (Bao et al., 2015), we detected that the expression of satellite RNAs and some human endogenous retroviruses (HERVs) were significantly increased upon depletion of TLKs (Figure 4B). Using RT-qPCR, we confirmed that the expression of several repetitive elements, including satellites and HERVs, were increased in U-2-OS cells, similar to TERRAs (Figure 3C and 4C). In contrast, other repetitive elements, such as 5S rDNA or Alu, were unaffected by TLK depletion (Figure 4C). ChIP-qPCR analysis showed that similar to telomeres (Figure 3B), there was a tendency towards decreased occupancy of H3.3 at HERV elements (Figure 4D), suggesting that reduced H3.3 may underlie desilencing upon TLK depletion.

**Figure 4.**
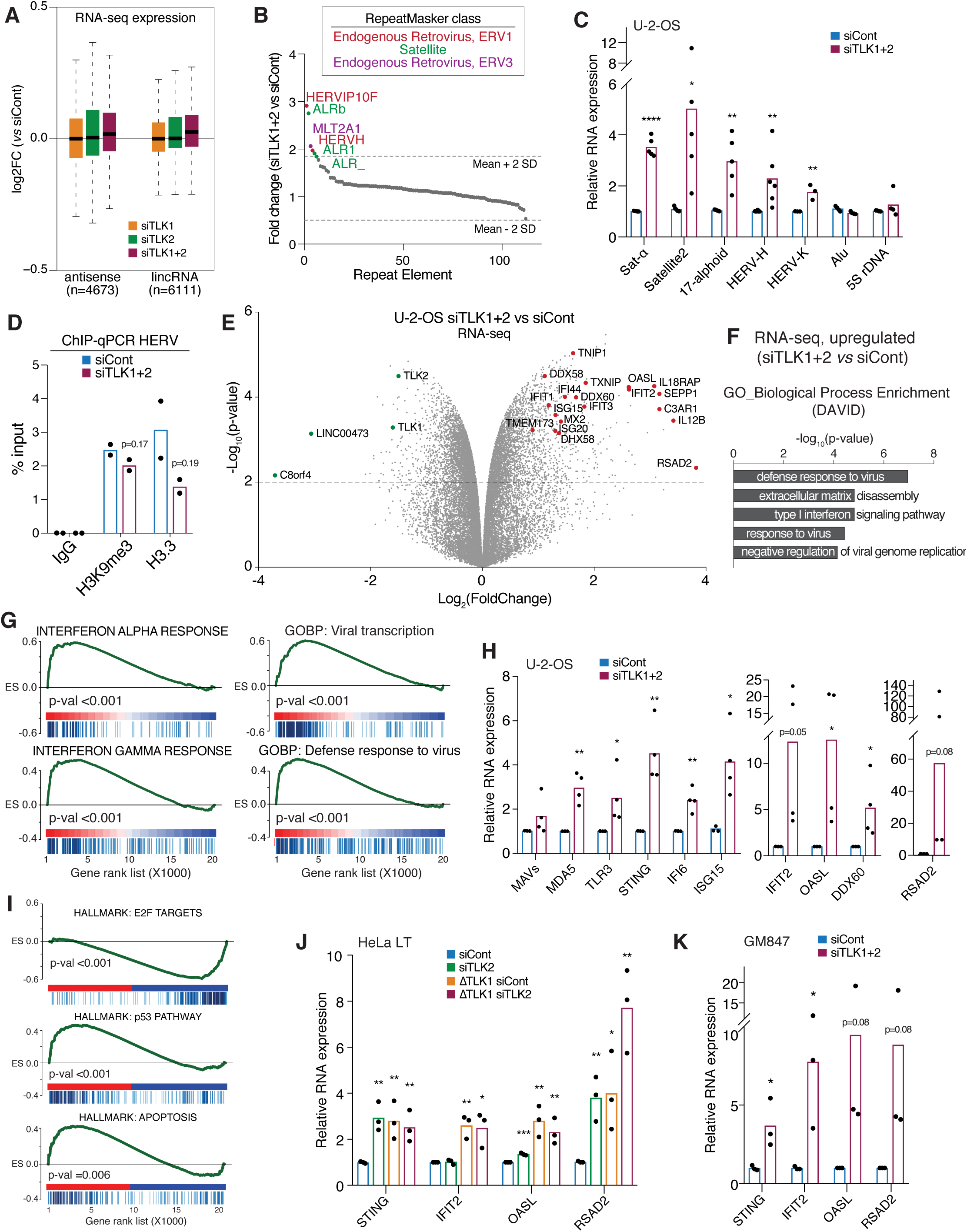
TLK depletion compromises silencing and activates innate immune signaling. **(A)** Boxplots of RNA-seq noncoding expression such as antisense RNA or lincRNA relative to siCont (n=2). **(B)** Analysis of repetitive RNAs expression (RNA-seq, n=2). Fold change in RNAs (siTLK1+2 vs siCont) transcribed from different repeat types were plotted as a rank order from highest to lowest. The horizontal dotted line represents a cut-off of 2 SD from the mean. Repeats enriched more than 2 SD from the mean are labelled, colours represent the RepeatMasker broad repeat class to which that repeat type belongs. **(C)** Expression of DNA repetitive elements by RT-qPCR in U-2-OS cells. Data were normalized to unchanging expression gene levels (B-actin) and the signal obtained in siCont conditions was set to 1 (n=5 Sat-*α*/Satellite2/17-alphoid, n=6 HERV-H, n=3 HERV-K, n=4 Alu/5S rDNA). **(D)** ChIP-qPCR was used to assess occupancy of H3.3 and H3K9me3 at HERV in U-2-OS cells. Data of qPCR amplification were normalized to input (n=2). **(E)** Volcano plot representing gene expression profile of siTLK1+2 versus siCont obtained from RNA-seq of U-2-OS cells (n=2). Grey dots indicate genes, green dots represent the most downregulated genes together with TLK1 and TLK2. Red dots represent genes belonging to the interferon response. P-values and fold changes are computed by fitting a linear model with the R package limma. **(F)** GO analysis (enrichment Biological Process) of RNA-seq differentially up-regulated genes upon double TLK knockdown (siTLK1+2). **(G)** GSEA of RNA-seq differentially expressed genes corresponding to the samples siTLK1+2 vs siCont. **(H)** Validation of expression levels of RNA and DNA sensors and interferon response genes by RT-qPCR in U-2-OS cells. Data were normalized to unchanging expression gene levels (B-actin) and the signal obtained in siCont conditions was set to 1 (n=4). **(I)** GSEA of RNA-seq differentially expressed genes corresponding to the samples siTLK1+2 vs siCont. **(J)** Expression levels of interferon response genes by RT-qPCR in HeLa LT cells, data were analysed as in (H) (n=3). **(K)** Expression levels of interferon response genes by RT-qPCR in GM847 cells, data were analysed as in (H) (n=3). ****P < 0.0001, ***P < 0.001, **P < 0.01, *P < 0.05, unpaired t test (Figure 4D), unpaired t test, one-tailed (Figure 4C, H, J-K)

In addition to genetic variants and repetitive regions, we analyzed differential gene expression in coding regions (−Log_2_ fold-change cut-off of 2). This analysis revealed that more genes were up-regulated than down-regulated (Figure 4E), with the strongest transcriptional change observed upon TLK loss being a potent innate immune response, comprised of the Type-I interferon (IFN) and TNF*α* programs. Gene ontology (GO) analysis confirmed that antiviral responses were among the categories most enriched in the upregulated genes upon total TLK loss (Figure 4F) and gene set enrichment analysis (GSEA) validated that IFN response gene expression signatures were upregulated in TLK depleted cells (Figure 4G). In addition, the checkpoint response (p53 pathway) and apoptosis signatures were also upregulated, whereas the E2F target signature was downregulated (Figure 4I), corroborating the p53 activation, G1/S cell cycle arrest and extensive cell death phenotypes we reported previously upon TLK depletion (Lee et al., 2018b). We further validated the activation of selected IFN response genes by RT-qPCR in U-2-OS cells, as well as in two additional cell lines, the ALT+ GM847 cell line and the ALT-HeLa LT cell line (Figure 4H, 4J and 4K). Notably, numerous genes involved in recognizing cytoplasmic or extracellular viral RNAs were activated, including *MDA5/IFIH1* and *MAVS/IPS-1*, involved in RNA sensing, as well as *STING/TMEM173*, an important mediator of IFN activation in response to cytosolic DNA/RNA detected by different sensors (Figure 4H and 4j-k), and accompanied by a number of other IFN-stimulated genes, including *IFI6, ISG15, IFIT2, OASL, DDX60* and *RSAD2*, that execute various antiviral functions (Figure 4H, 4J and 4K).

### TLK depletion upregulates IFN independently of replication stress signaling and ALT

The activation of cGAS-STING driven immune signaling has been clearly linked with the maintenance of genome stability (Bakhoum et al., 2018; Chen et al., 2017; Harding et al., 2017; Li and Chen, 2018; Mackenzie et al., 2016; Mackenzie et al., 2017). In particular, ALT+ tumors exhibit epigenetically silenced STING, presumably to prevent its activation by cGAS-dependent detection of extrachromosomal telomeric DNA (Chen et al., 2017; Wu et al., 2018), and SAMHD1 has been demonstrated to prevent the IFN response during replication stress through the recruitment of the MRE11 nuclease (Coquel et al., 2018). To investigate the relationship between replication stress and the innate immune response following TLK depletion, we examined the requirements for upregulation of *RSAD2* and *STING*. Depletion of STING in HeLa LT cells severely abolished *RSAD2* gene induction while not influencing replication stress signaling (Figure 5A and 5B). Canonical STING activation induces type-I IFN genes through the IRF3 transcription factor and requires the activity of the TANK-binding kinase 1 (TBK1) (Chen et al., 2016). Treatment of cells with the TBK1 inhibitor BX795 completely abolished the upregulation of *RSAD2*, but not *STING*, following TLK depletion (Figure 5C).

**Figure 5.**
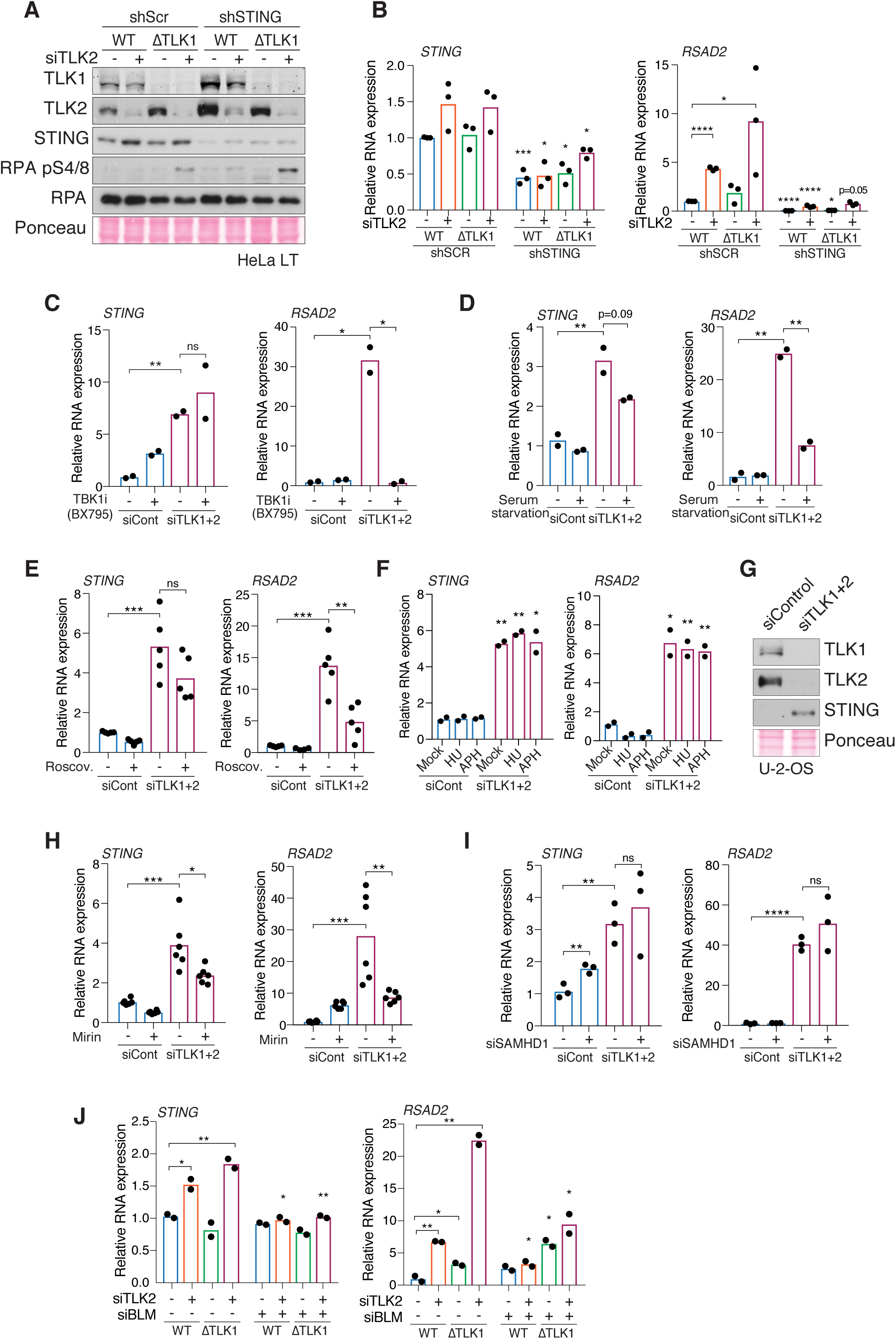
TLK inhibition upregulates innate immunity independently of replication stress signaling. **(A)** Western blot of HeLa LT cells infected with scramble shRNA or shRNA against STING, 48 h after being treated with the corresponding siRNAs. **(B)** Expression levels of STING and RSAD2 by RT-qPCR in HeLa LT cells 48 h after being treated with the corresponding siRNAs. Data were normalized to unchanging expression gene levels (B-actin) and the signal obtained in siCont conditions was set to 1 (n=3). **(C)** Expression levels of STING and RSAD2 by RT-qPCR in U-2-OS cells mock treated or treated with 1 µM TBK1 inhibitor BX795 for 24 h. Data were analysed as in (B) (n=2). **(D)** Expression levels of STING and RSAD2 by RT-qPCR in control or serum starved U-2-OS cells. Data were analysed as in (B) (n=2). **(E)** Expression levels of STING and RSAD2 by RT-qPCR in U-2-OS cells mock treated or treated with 50 µM Roscovitine for 5 h. Data were analysed as in (B) (n=5). **(F)** Expression levels of STING and RSAD2 by RT-qPCR in U-2-OS cells mock treated or treated with either 0.5 mM HU for 2 h or 0.5 µM Aphidicolin for 4 h. Data were analysed as in (n=2). **(G)** Western blot of STING in U-2-OS cells, 48 h after being treated with the corresponding siRNAs. **(H)** Expression levels of STING and RSAD2 by RT-qPCR in U-2-OS cells mock treated or treated with 50 µM Mirin for 5 h. Data were analysed as in (B) (n=6). **(I)** Expression levels of STING and RSAD2 by RT-qPCR in U-2-OS cells 48 h after being treated with the corresponding siRNAs. Data were analysed as in (B) (n=3). **(J)** Expression levels of STING and RSAD2 by RT-qPCR in HeLa LT cells 48 h after being treated with the corresponding siRNAs. Data were analysed as in (B) (n=2). ****P < 0.0001, ***P < 0.001, **P < 0.01, *P < 0.05, unpaired t test (Figure 5B-F, H-J)

We previously demonstrated that replication stress signaling in TLK depleted cells was dependent upon active DNA replication (Lee et al., 2018b). Similarly, we found that the activation of innate immune signaling upon TLK loss was strongly abrogated following serum starvation or treatment of cells with the CDK2 inhibitor Roscovitine (Figure 5D and 5E). Conversely, treatment of U-2-OS cells with HU and APH to dramatically upregulate replication stress signaling (Figure 1) did not activate *RSAD2* or *STING* expression (Figure 5F). In contrast, TLK depletion increased STING mRNA and protein levels and this effect was not enhanced by HU or APH treatment (Figure 5F, 5G, S4A and S4B).

To further examine the potential links between replication forks and innate immune signaling in TLK depleted cells, we examined the impact of inhibiting of depleting several replication fork response proteins. Treatment with the MRE11 inhibitor Mirin attenuated replication stress signaling (Figure 1F) and also alleviated *RSAD2*, and to a lesser extent, *STING* induction, indicating that MRE11 activity was also required for full induction of innate immunity following TLK depletion (Figure 5H). Consistent with recent reports, depletion of SAMHD1 mitigated p-RPA signaling in HU treated cells (Figure S4C) (Coquel et al., 2018). However, it did not impair p-RPA signaling in HU treated cells depleted for TLK1/2 (Figure S4C) and was dispensable for the induction of both *RSAD2* and *STING* following TLK depletion (Figure 5I). We also examined the influence of BLM depletion, which reduced replication stress signaling and inhibited C-circle production (Figure 1G, 1H, 3N and S3F). Depletion of BLM attenuated the activation of both *STING* and *RSAD2* in HeLa LT cells following TLK depletion (Figure 5J).

Together, our results indicated that depleting the TLKs results in the induction of a canonical STING-TBK1-dependent IFN response, even in cells where STING has been epigenetically silenced. This response requires DNA replication, reflecting the fact that TLK depletion phenotypes require passage through S-phase where histone exchange levels are their highest, but is not influenced by the intensity of replication stress signaling per se. The induction of the IFN response in TLK depleted cells is linked to C-circle production, since it is BLM dependent. Thus, in the context of cancer cells with a dynamic chromatin state, the effects of faulty heterochromatin maintenance observed in TLK depleted cells account for induction of innate immunity.

### TLK expression correlates with suppressed innate immune signaling in human cancer

Tumors with unstable genomes, micronuclei and ALT+ telomeres generate sources of immunostimulation that can activate STING and induce an IFN response if detected by cytosolic sensors (Bakhoum and Cantley, 2018; Chen et al., 2017). As the IFN response in tumor cells may recruit cytotoxic immune cells and promote tumor cell apoptosis, many tumors circumvent this pathway by silencing STING epigenetically to minimize immune surveillance (Chen et al., 2017; Wu et al., 2018), Thus, we next wanted to determine if our conclusions from tissue culture cells were evident in data from human tumor patients. We generated an expression signature of the 10 most highly upregulated IFN genes belonging to the GSEA IFN Alpha/Gamma signatures following TLK depletion (Figure 4G and 6A) and analyzed publically available datasets from The Cancer Genome Atlas (TCGA) for correlations with the levels of *TLK1* and *TLK2* across different tumor types. Expression levels of *TLK1* and *TLK2* anti-correlated significantly with those of STING (encoded by the *TMEM173* gene), and this occurred in most tumor types analyzed (Fig 6B and 6C). The analysis also indicated that *TLK1* and *TLK2* expression levels positively correlate in most tumor types analyzed (Figure 6C). In contrast, TLK expression anti-correlated with our 10 gene IFN signature (Figure 6A), TNF*α* GSEA signature and several TNF*α* genes in several tumor types (Figure 6C and 6D). We reasoned that tumors with high levels of genome instability may select both for higher levels of TLK activity to suppress toxic replication stress and for STING silencing. We examined chromosomal instability (CIN) levels in tumors by using a previously defined signature containing a 25-gene set (Carter et al., 2006), as well as the aneuploidy score estimated from CGH SNP arrays, so it is a measure independent from expression (Taylor et al., 2018). While *TLK1* expression levels only positively correlated with CIN/aneuploidy levels in selected tumor types (Figure 6C), *TLK2* expression significantly correlated with CIN/aneuploidy levels in most tumor types (Figure 6C). Following these correlations, we next addressed whether TLK^high^-STING^low^-CIN^high^ tumors were differentially infiltrated by various immune cell populations. We took measurements of leukocyte and stromal fractions based on methylation and expression signatures (Taylor et al., 2018), as well as immune cell signatures of populations responding to innate immune signaling to detect tumor cells: cytotoxic CD8+ T cells, activated NK cells and M1 macrophages (Thorsson et al., 2018). We detected several tumor types with significantly reduced fractions of immune infiltration correlating with high TLK expression (Figure 6C). Many of these presented as well with TLK2 levels positively correlating with CIN/aneuploidy and negatively correlating with STING. Interestingly, some of the more significant tumor types with this behavior are described to be immune infiltration depleted or “cold” tumors such as ACC, LIHC, GBM, SARC, LGG (Figure 6C) (Thorsson et al., 2018). Furthermore, several of these tumor types are also frequently ALT+ (Ceccarelli et al., 2016; Lee et al., 2018a; Zheng et al., 2016).

**Figure 6.**
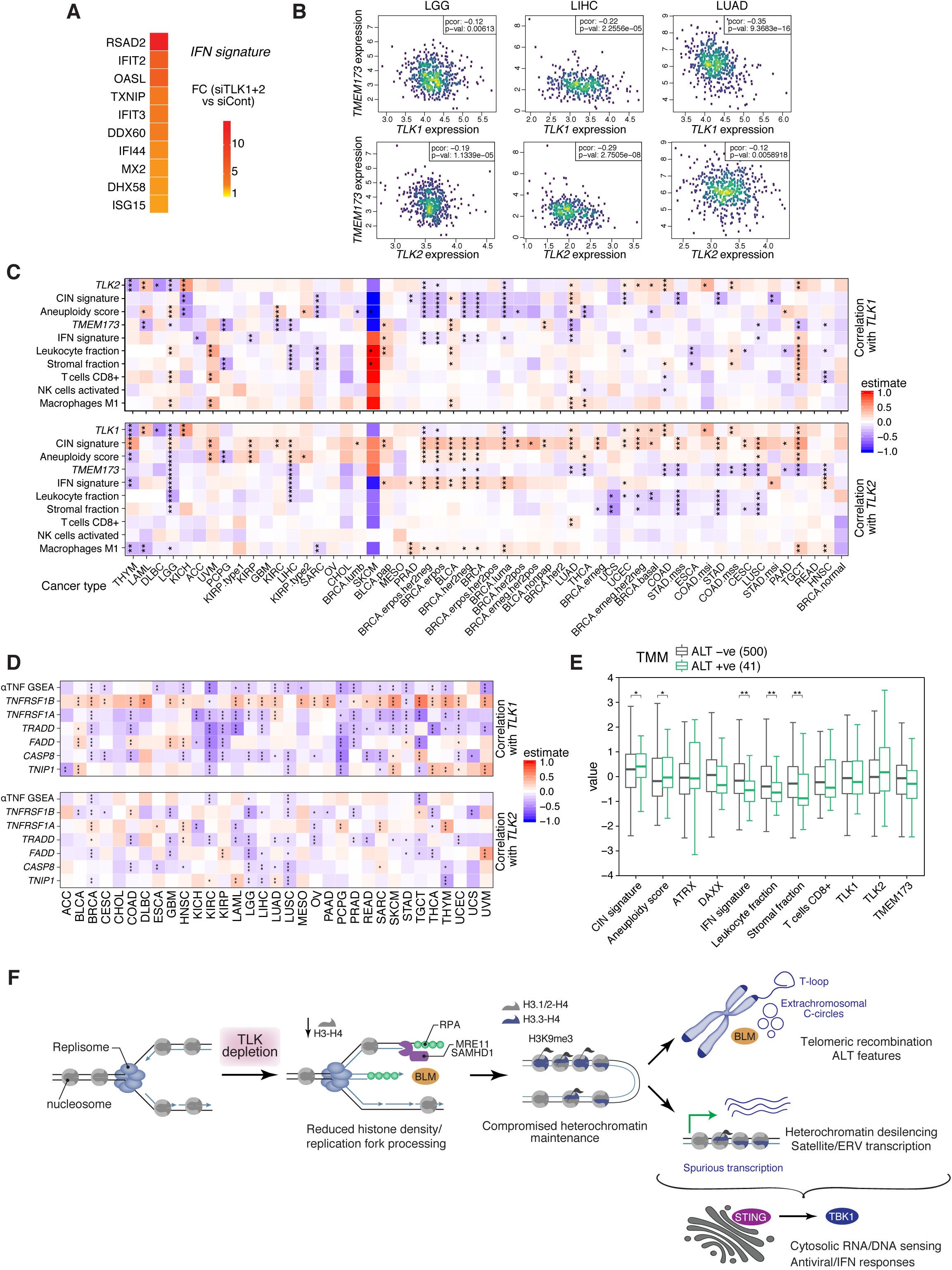
TLK expression is associated with enhanced CIN and suppressed innate immune signalling in human cancer. **(A)** Fold changes of expression levels of the 10-gene IFN signature generated from our RNA-seq results in U-2-OS cells. **(B)** Correlation between *TLK1/TLK2* and STING (*TMEM173*) in the TCGA datasets of Low Grade Glioma (LGG), Liver Hepatocellular Carcinoma (LIHC) and Lung Adenocarcinoma (LUAD). **(C)** Heatmaps with correlations of expression levels of *TLK1* and *TLK2* with CIN signature, Aneuploidy score, STING (*TMEM173*), the IFN signature defined in (A), and different signatures of tumour immune populations across different TCGA datasets. **(D)** Heatmaps with correlations of expression levels of *TLK1* and *TLK2* with TNF*α* signature as well as individual TNF pathway genes. **(E)** Boxplot of expression levels of genes, CIN signature, Aneuploidy score, the IFN signature defined in (A), and different signatures of tumour immune populations. The analysis was performed in different merged TCGA datasets where TMM status is defined. **(F)** Model on how TLK activity maintains heterochromatin state and the loss of TLK activity promotes telomere recombination and spurious transcription, triggering an innate immune response.

Last, we interrogated whether tumors in the TCGA with different telomere maintenance mechanism (TMM) would select to express differentially the signatures described above. Overall, we found a positive correlation between ALT+ tumors and CIN/aneuploidy levels, whereas ALT+ tumors anti-correlated with our IFN signature and signatures of immune infiltration (Figure 6E). Despite not significant due to variability across cancer types and low N, ALT+ tumors showed a tendency towards higher levels of TLK2 and lower levels of STING.

These data indicate that tumors that harbor high levels of genome instability (CIN^high^ or ALT+ tumors) may select for high TLK levels and defective cytosolic DNA sensing by STING silencing, as means to prevent tumor-infiltrating lymphocytes. Therefore, these TLK^high^-STING^low^-CIN^high^ or TLK^high^-STING^low^-ALT+ tumors may benefit from inhibition of TLK activity, promoting immune detection and suppressing cancer cell survival.

## Discussion

Here, we further establish that TLKs play a critical role in chromatin maintenance that protects replication forks and demonstrate that heterochromatin maintenance at telomeres and other repetitive genome elements is sensitive to TLK depletion. TLK depletion impairs histone deposition, including that of the H3.3 variant, and leads to epigenome instability characterized by desilencing of endogenous viral elements and ncRNAs, enhanced features of ALT and re-activation of STING-mediated immune responses (Figure 6F). These effects vary between cell lines, but were the most dramatic in ALT+ U-2-OS cells, where loss of TLK activity also led to increased chromatin accessibility at heterochromatic loci, loss of HP1α and spurious transcription of non-coding RNA. Our findings also indicate that CIN^high^ tumors may select for high levels of TLK expression and that targeting TLK activity activates the IFN response, even in cases where it has been epigenetically silenced. STING agonists have shown promise in preclinical studies, as stand-alone agents or in enhancing the effects of immunotherapy, but the interferon response has also been implicated in suppressing immunotherapy and shown to promote metastasis in CIN^high^ tumors (Abramov et al., 2010; Bakhoum et al., 2018; Benci et al., 2016; Curran et al., 2016). Thus, it is likely that important context-dependent effects of agents that activate STING will need to be considered.

Activation of cGAS-STING mediated innate immunity by DNA damage and replication stress through the recognition of micronuclei has been well documented (Harding et al., 2017; Mackenzie et al., 2017). Recent work has also shown that SAMHD1 suppresses IFN response activation following treatment with HU in a manner dependent on the activity of the MRE11 nuclease, thus implicating byproducts of replication fork processing (Coquel et al., 2018). We identified a clear separation of function, as SAMHD1 is required for replication stress signaling but dispensable for innate immune activation following TLK depletion. Even after HU induction, that strongly increases DDR signaling, there was no correlation between ssDNA or RPA phosphorylation and IFN activation (Figure 5I and S4C). Thus, TLK depletion, despite causing less damage than HU or APH, is able to more robustly trigger the IFN response. Perturbation of the epigenetic landscape by depleting H3.3, multiple H1 genes or DNA methylases, resulted in transcription of endogenous RNA from heterochromatin, including HERVs and other non-coding repeat elements, similar to what we have observed here following TLK depletion (Chiappinelli et al., 2015; Elsasser et al., 2015; Izquierdo-Bouldstridge et al., 2017). Moreover, desilencing of endogenous repeats was accompanied by the activation of an antiviral response in some of these cases (Chiappinelli et al., 2015; Izquierdo-Bouldstridge et al., 2017). TLK activity has not previously been implicated in the silencing of endogenous viruses, however TLK2 was identified in a screen for the latency of gammaherpesvirus, that is known to be maintained through an epigenetic mechanism (Dillon et al., 2013). Thus, the impact of TLK depletion on innate immune signaling is potentially due to its combined roles in preventing DNA damage and maintaining the epigenetic landscape and it is likely that multiple sensors that activate STING are involved (Chen et al., 2016).

ALT+ cells are typically characterized by high levels of CIN, higher basal levels of replication stress and display complex karyotypes (Gagos et al., 2008;Marzec et al., 2015). In addition, the C-circles generated by ALT induce the cGAS-STING pathway and have been proposed to provide selective pressure for the epigenetic silencing of STING in order to suppress potentially cytostatic innate immune signaling (Chen et al., 2017). ALT+ cells may also have a higher dependency on ASF1, and thus TLK activity, to support H3.3 deposition and heterochromatin maintenance in the absence of functional ATRX-DAXX (Lovejoy et al., 2012). This is consistent with previous work that has shown redundancy in the histone chaperone network that can mitigate deleterious effects arising upon histone pool or histone chaperone imbalances (Lacoste et al., 2014;Drane et al., 2010). Although TLK activity suppresses ALT, TLK dysfunction is unlikely to be an ALT driver event in cancer, since both TLK1 and TLK2 are very rarely mutated or deleted in cancer genomes and are required for DNA replication (Lee et al., 2018b). Instead, ALT+ cells may be addicted to TLK activity in order to support heterochromatin maintenance in the absence of ATRX-DAXX and to circumvent innate immune responses (Chen et al., 2017). Our results therefore suggest that reactivation of STING expression in ALT+ cancer cells upon TLK inhibition would be a potential strategy to re-sensitize ALT+ cells to C-circles and other cytoplasmic DNA fragments. This inflammatory response may render cancer cells vulnerable to cytotoxic immune cell recruitment and enhance immunotherapy approaches. Recent work reporting inhibitors for CDK9, a key regulator of transcription and epigenome maintenance, supports this possibility (Zhang et al., 2018).

Taken together, we have uncovered a complex connection between TLK-mediated chromatin maintenance, genomic stability and the innate immune response. These results suggest that TLK activity promotes immune suppression in CIN^high^ tumors, indicating that its targeting could induce IFN signaling regardless of telomere status or epigenetic silencing of effector genes. TLK inhibitors could therefore represent a novel rational targeted therapy to render ALT+ or CIN^high^ cancers more vulnerable to the induction of cell death and provide a window of opportunity to reset chromatin state, enhance existing therapies and provoke tumor regression. Moreover, the newly uncovered role of TLK activity in the suppression of innate immune mediated inflammation may be highly relevant to the etiology of intellectual disability/autism spectrum disorder in patients with germline *TLK2* mutations (Lelieveld et al., 2016; Reijnders et al., 2018).

## Materials and Methods

### Cell lines, growth conditions and drug treatments

The human cell lines U-2-OS, HeLa LT, GM847, RPE-1, AD293 and HEK-293 were grown in DMEM (Thermo Fisher Scientific) supplemented with 10% FBS (Sigma-Aldrich) and penicillin–streptomycin at 37°C in a 5% CO2 incubator. Knockout ΔTLK1 U-2-OS and HeLa LT cell lines were generated using CRISPR/Cas9 genome editing. Guide RNA sequence targeting human TLK1 exon 10 (TAACTGTTGTAAAGTGCCCG) were cloned into the plasmid pX330-CRISPR-Cas9-SV40prom-EGFP (Cong et al., 2013) after digestion with BbsI (NEB). Cells were transfected using Lipofectamine 2000 (Thermo Fisher Scientific) with the corresponding pX330-CRISPR-Cas9-SV40prom-EGFP plasmids and 48 hours following transfection, single cells positive for GFP were FACS-sorted (BD FACSAria III) at a ratio of 1 cell per well in 96 well plates. Single cell clones were expanded and screened by Western Blot for protein levels of TLK1. For lentiviral transductions of shRNA against STING, AD293 producer cells were transfected using PEI (Polysciences Inc.) with 10 μg of pLKO.1-puro-Scramble control (SHC002) or pLKO.1-puro-shSTING (TRCN0000163296) and with lentiviral packaging vectors (2 μg REV, 6 μg RRE and 2 μg VSV-G). Medium was refreshed 8 h after transfection and viral supernatants were collected and filtered through a 0.45 μm filter at approximately 48 and 72 h post-transfection. For infection, cells were overlaid with filtered viral supernatant supplemented with 8 μg/ml Polybrene (Sigma-Aldrich) twice to achieve optimal infection efficiency. Cells were selected with 1 μg/ml of puromycin.

For drug treatment we used: Hydroxyurea (Sigma), Aphidicolin (Calbiochem) at the indicated doses; Roscovitine (Tocris), 50 μM for 5 hours; Mirin (Tocris), 50 μM for 5 hours; BX795 (Selleckchem), 1 μM for 24 hours; DRB (Sigma-Aldrich), 100 µM for 3 h. For serum starvation experiments, after cell attachment the medium was replaced with low-serum media (0.25% FBS) for serum-starved cells and with fresh media (10% FBS) for asynchronous cells. After 24 h, medium was replaced again (0.25% FBS for serum starved cells; 10% FBS for control cells) until cells were harvested.

### siRNA transfections

For siRNA transfection, cells were transfected with siRNAs at a final concentration of 100 nM (Sigma-Aldrich) using Lipofectamine RNAiMAX (Thermo Fisher Scientific). Transfections were done on attached cells that had been plated for approximately 18 hours, in media without antibiotics and the transfection mix was prepared in Opti-MEM (Thermo Fisher Scientific). Cells were transfected once and either split for several experiments after 24 hours and harvested after additional 24-48 hours or directly harvested 48 hours post-transfection. We used luciferase and GFP targeting as mock negative controls (siGL2: 5’-CGUACGCGGAAUACUUCGA-3’ and siGFP: 5’-GGCUACGUCCAGGAGCGCCGCACC-3’), as well as ON-TARGETplus Non-targeting Control Pool (Dharmacon D-001810-10). The sequences of the forward siRNAs used in this study are the following: siTLK1 (5’-GAAGCUCGGUCUAUUGUAA-3’), siTLK1 (Dharmacon, SMARTpool ON-TARGETplus L-004174-00), siTLK2#1 (5’-GGAGGGAAGAAUAGAUGAU-3’), siTLK2#2 (5’-GGAAAGGAUAAAUUCACAG-3’), siTLK2 (Dharmacon, SMARTpool ON-TARGETplus L-005389-00), siASF1a (5’-AAGUGAAGAAUACGAUCAAGU-3’), siASF1b (5’-AACAACGAGUACCUCAACCCU-3’), siSAMHD1 (5’-CUCGUCCGAAUCAUUGAUACA-3’), siMRE11 (5’-GCUAAUGACUCUGAUGAUA-3’), siCTIP (5’-GCUAAAACAGGAACGAAUC-3’), siBLM (Dharmacon, SMARTpool ON-TARGETplus L-007287-00).

### Clonogenic cell survival assays

For survival analysis upon drug treatments, siRNA-transfected U-2-OS cells were seeded onto twelve-well plates in technical duplicates and after 24 hours of plating, cells were treated with Hydroxyurea or Aphidicolin at the indicated doses for 24 hours and then washed and grown in fresh medium. Cells were incubated at 37°C for 15 days and were then fixed and stained with crystal violet (Sigma-Aldrich). Colonies were counted using an in-house–built ImageJ macro using a Trainable Weka Segmentation plugin. On the basis of the colony number, plating efficiency (PE = number of colonies formed/number of cells seeded) and surviving fraction (SF = number of colonies formed after drug treatment/number of cells seeded × PE) were calculated and plotted.

### Western blot analysis

Cells were collected after two cold PBS washes by scraping in 2X SDS Lysis Buffer (4% SDS, 20% Glycerol, 120mM Tris-Cl pH 6.8, 1x protease (Roche) and phosphatase inhibitors (Sigma-Aldrich)) on ice. Lysates were sonicated at medium-high intensity for 10 minutes in a Bioruptor Standard (Diagenode) placed at 4°C and subsequently boiled for 10 minutes at 90°C. Proteins were quantified using the DC Protein Assay (Bio-Rad), separated by SDS-PAGE and transferred to 0.2 µm pore Nitrocellulose membrane (Amersham Protran; Sigma-Aldrich). Membranes blocked with 5% milk/PBST for 1 h at room temperature and probed with primary antibodies overnight at 4°C. Primary antibodies used were against the following: TLK1 (Cell Signaling #4125, 1:1000); TLK2 (Bethyl Laboratories A301-257A, 1:1000); RPA2 pS4/S8 (Bethyl Laboratories A300-245A, 1:500); RPA2 pS33 (Bethyl Laboratories A300-246A, 1:500); RPA2 (Calbiochem NA19L, clone RPA34-20, 1:1000); H2AX pS139 (γH2AX) (Santa Cruz sc-101696, 1:500; Cell Signaling #2577 1:1000); p53 pS15 (Cell Signaling #9284, 1:500); MRE11 (Santa Cruz sc-135992, 1:200); CtIP (Santa Cruz sc-271339, 1:500); BLM (Abcam ab2179, 1:2000); STING (R&D Systems MAB7169, clone 723505, 1:1000). These were detected with appropriate secondary antibodies conjugated to HRP and visualized by ECL-Plus (GE Healthcare).

### Immunofluorescence (IF), immuno-FISH and high-throughput microscopy (HTM)

For IF of chromatin-bound proteins, cells were grown on 8-well Lab Tek II chamber slides (Labclinics) to sub-confluence. For regular IF, cells were grown on Poly-L-Lysine coated coverslips. For chromatin-bound proteins, pre-extraction was performed using cold 0.2% Triton X-100 in 1X PBS on ice for 5 minutes. Cells were fixed for 10 minutes in 4% paraformaldehyde at room temperature and permeabilized for 5 minutes in 0.2% Triton in 1X PBS at room temperature. After two washes in PBS, fixed cells were incubated for 1 h in blocking solution (3% BSA 0.1%Tween/PBS) and stained using the primary antibodies indicated below diluted 1:250 in blocking solution for 4 h at room temperature in a humid chamber. Primary antibodies used were the following: RPA2 (Calbiochem NA19L, clone RPA34-20); H2AX pS139 (γH2AX) (Santa Cruz sc-101696; Cell Signaling #2577); HP1α (Active Motif 39977, clone 2HP1H5,): PML (Santa Cruz sc-966, clone PG-M3); TRF2 (Novus Biologicals NB110-57130). The secondary antibodies used were Alexa Fluor 488 (goat anti-rabbit IgG), Alexa Fluor 488 (goat anti-mouse IgG), Alexa Fluor 568 (goat anti-rabbit IgG), and Alexa Fluor 568 (goat anti-mouse IgG) from Thermo Fisher Scientific, and were used at a dilution of 1:500 in blocking solution. Slides were mounted in Vectashield (Vector Labs).

For IF-FISH, after standard IF staining with primary and secondary antibody incubations, washes were performed with IF blocking solution. To continue with FISH, slides were fixed again in 4% formaldehyde for 5 min at room temperature, washed in PBS and dehydrated in increasing concentrations of ethanol (70%, 95%, and 100%). The slides were air-dried, incubated with Telomere PNA Probe/Cy3 (Dako-Agilent) and denatured on an 80°C hot plate for 5 minutes. Hybridization took place in the dark for 3-4 hours at room temperature. The slides were subsequently washed in Rinse Solution (Dako-Agilent), washed in pre-heated Wash Solution (Dako-Agilent) and dehydrated in increasing concentrations of ethanol (70%, 95%, and 100%). Slides were washed in DAPI and mounted in Vectashield (Vector Labs).

Confocal fluorescence images were obtained on a Leica DM2500 SPE confocal system. Images were taken with 40x NA 1.15 oil or 63x NA 1.3 oil objectives and the standard LAS-AF software. For high-throughput microscopy (HTM), 24-48 images were automatically acquired from each well with a robotized fluorescence microscopy station (Scan^R; Olympus) at 40× magnification and non-saturating conditions. Images were segmented using the DAPI staining to generate masks matching cell nuclei from which the corresponding signals were calculated using an in-house–developed package based on Cell Profiler.

### RNA isolation and quantitative real time PCR (qRT-PCR)

Cells were collected after two cold PBS washes by scraping in Tri-Reagent (Sigma-Aldrich). RNA was isolated by chloroform extraction followed by centrifugation, isopropanol precipitation, washing two times in 75% ethanol and resuspension in DEPC-treated water. Nucleic acid quantification was performed with a Nanodrop 8000 Instrument (Thermo Fisher Scientific). Reverse transcription reaction was carried out using High Capacity cDNA Reverse Transcription Kit (ABI), following the manufacturer’s instructions, in a reaction volume of 20 μl and with Random primers contained in the kit. cDNA was stored at −20°C. For the detection on Telomeric repeat-containing RNA (TERRA), a first strand cDNA synthesis was first performed with the TERRA-specific RT-primer 5’-CCCTAACCCTAACCCTAACCCTAACCCTAA-3’ (Feretzaki and Lingner, 2017) and a housekeeping gene-specific primer (Reverse) was included in the same reaction for normalization purposes. The reaction was carried out at 55°C in the presence of RNA inhibitor SUPERase IN (Thermo Fisher Scientific) using the SuperScript III RT (Thermo Fisher Scientific). qPCR was performed using the comparative CT method and a Step-One-Plus Real-Time PCR Instrument (Applied Biosystems). SYBR Green reactions were carried out in technical duplicate or triplicate in a final volume of 10 μl. For SYBR Green 1X SYBR Green PCR Master Mix (no. 4364344; ABI) or 1X SYBR Select Master Mix (no. 4472908; ABI), forward and reverse primer (Sigma; 100-500 nM each) and 25 ng of template were used. Thermocycling parameters used were: 95 °C 20 s; 40 cycles 95 °C 3 s, 60 °C 30 s; melting curve. The 2−ddCT method was used for the analysis of the amplification products. Primer pairs used are indicated in Table 1.

**Table 1.**
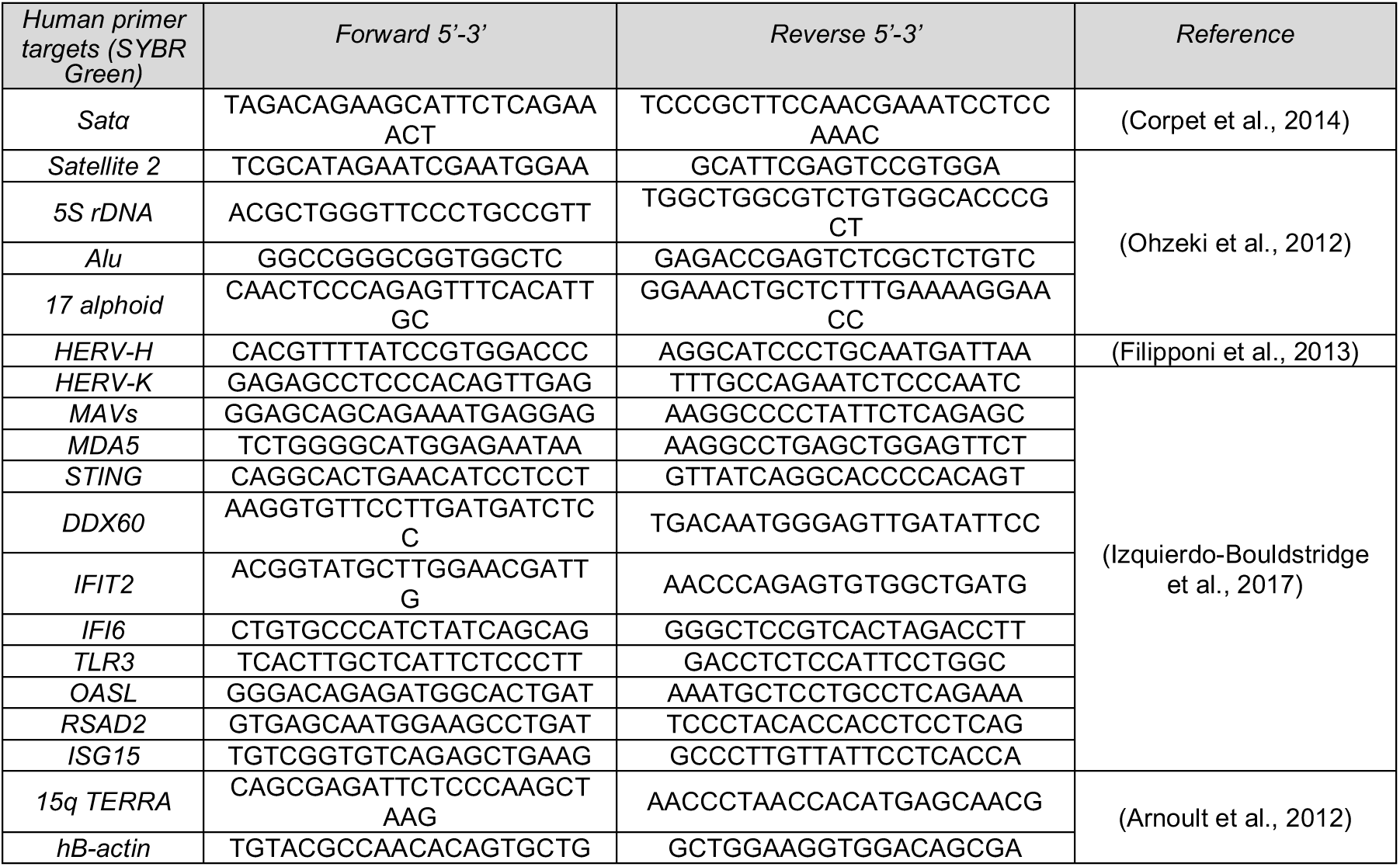
Primer pairs used for qRT-PCR.

### Assay for Transposase-Accessible Chromatin (ATAC)-seq and -qPCR

The ATAC protocol was adapted from (Buenrostro et al., 2013). U-2-OS or HeLa LT cells were transfected in biological duplicate with siRNAs (siCont, siTLK1, siTLK2, siTLK1+2 for U-2-OS, and siCont, siTLK2 for HeLa LT WT and ΔTLK1) and 48 hours post-transfection were collected by trypsinization and counted. 50 000 cells for U-2-OS or 75 000 cells for HeLa LT were resuspended in 50 μl of cold lysis buffer (10 mM Tris-HCl, pH 7.4, 10 mM NaCl, 3 mM MgCl2, 0.1% (v/v) Igepal CA-630) for nuclei preparation and treated with Nextera Tn5 Transposase (Nextera DNA Library Preparation Kit, Illumina) at 37°C for 30 min. DNA was first purified using MinElute PCR Purification Kit (Qiagen). Samples were amplified by PCR using NEBNext High-Fidelity 2x PCR Master Mix (NEB) using PCR primers Ad1_noMX and Ad2_Barcode (Sigma-Aldrich, see Table 2, (Buenrostro et al., 2013) for 5 initial cycles (1 cycle: 5 min 72°C, 30 sec 98°C; 5 cycles: 10 sec 98°C, 30 sec 63°C, 1 min 72°C). A qPCR side reaction was run on a tenth of the previous 5-cycle sample in order to determine the appropriate number of PCR cycles (N) and be able to stop amplification prior to saturation, in the presence of SYBR Green in an Applied Biosystems 7900HT Fast Real-Time PCR System (1 cycle: 30 sec 98°C; 20 cycles: 10 sec 98°C, 30 sec 63°C, 1 min 72°C). Then, the 5-cycle sample was run for an additional N=6 cycles, so the library had been amplified for a total of 11 cycles. Amplified libraries were purified using PureLink PCR Purification Kit (Thermo Fisher Scientific). The quality of purified libraries was assessed using a Bioanalyzer High-Sensitivity DNA Analysis kit (Agilent). For ATAC-seq, an equimolar pool was generated and the pool was sequenced in 2 lanes 50 nt paired-end in an Illumina HiSeq2500 (Illumina). ATAC-qPCR was performed with the same primers as in the ChIP-qPCR experiments (Table 3) on a 1:20 dilution of the eluted DNA after library amplification.

**Table 2:**
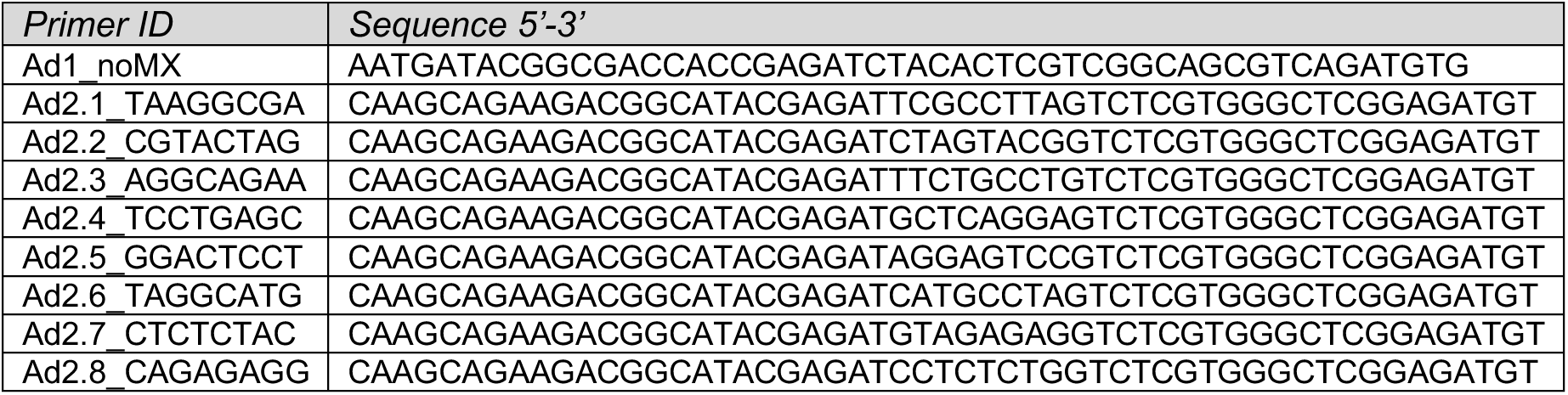
Primers used for ATAC library amplification

**Table 3:**
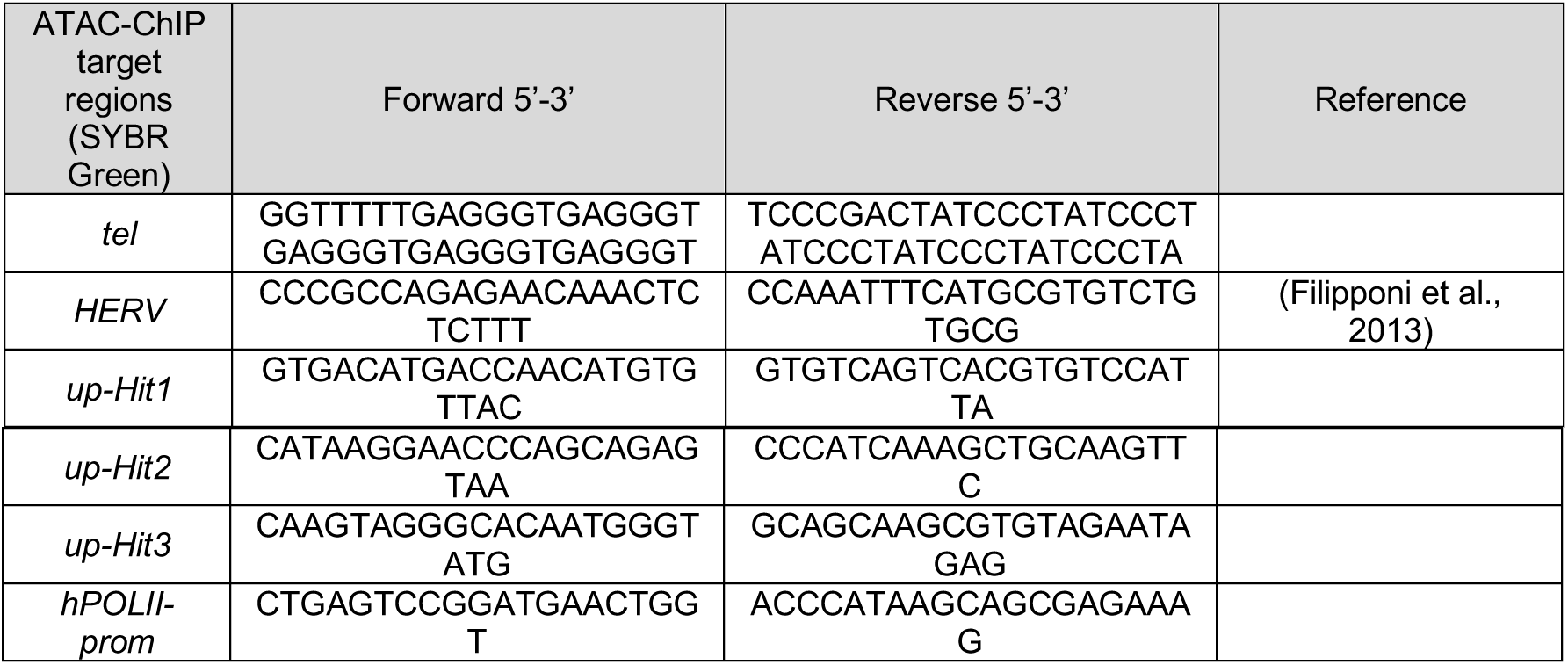
Primers used for ATAC-qPCR or ChIP-qPCR assays

For ATAC-seq analysis, paired end reads of length 50 bp were aligned to the human genome version hg19 using bowtie v0.12.9 (Langmead et al., 2009) with default parameters except for n=1 to limit the maximum number of mismatches in the seed and m=1 to report the best possible alignment for a given read. Alignments were sorted and indexed using Sambamba v0.5.8 (http://lomereiter.github.io/sambamba/). Duplicated reads were removed using Sambamba. Peaks were called with MACS v1.4.2 (Zhang et al., 2008) for each sample without any control. All parameters were left as default except for read length. Peaks were annotated with the HOMER v2.8.2 (Heinz et al., 2010) software with the hg19 annotations. Consensus peaks were defined as the union of all peaks from all samples. For each sample, the number of reads per peak was computed using the countBam function from the Rsamtools R package (version 1.30.0, http://bioconductor.org/packages/release/bioc/html/Rsamtools.htm). Differential accessibility was computed using DESeq2 (Love et al., 2014) with replicate as covariate. For ATAC-seq overlap with epigenetic marks, epigenetic information was downloaded from the Encode website (Consortium, 2012) (H3K9me3 [ENCFF001VDL]). Consensus peaks were classified according to their overlap with each epigenetic track [minimum overlap length 100bp]. P-values were computed with a Mann-Whitney test as implemented in the function wilcox.test in R (R Development Core Team, 2016). For ATAC-seq FC correlation between replication time and methylation marks, the Repliseq track (wgEncodeUwRepliSeqNhekWaveSignalRep1.bigWig) was downloaded from the Encode website in Bigwig format. H3K36me3 and H3K9me3 corresponding to the U-2-OS cell line were also downloaded in Bigwig format from the same repository. Mean normalized signal was computed for 10kb bins along the genome. Spearman correlations coefficients were computed with the cor function in R. For Figure S2C, Repliseq scores were binned in 25 quantiles. For ATAC-seq FC overlap with colors of chromatin, tracks were downloaded from Encode (wgEncodeAwgSegmentation ChromhmmHelas3.bed) corresponding to the ChromHMM algorithm (Ernst and Kellis, 2012) in the HeLa-S3 cell line. Peaks were classified to the color with larger overlapping segment. For better visualization, colors were collapsed to 10 classes as follows: Active Promoter = {Tss, TssF}, Promoter Flanking = {PromF}, Inactive Promoter = {PromP}, Candidate Strong enhancer = {Enh, EnhF}, Candidate Weak enhancer/DNase = {EnhWF, EnhW, DnaseU, DnaseD, FaireW}, Distal CTCF/CandidateInsulator = {CtcfO, Ctcf}, Transcription associated={Gen5, Elon, ElonW, Gen3, Pol2, H4K20}, Low activity proximal to active states= {Low}, Polycomb repressed = {ReprD, Repr, ReprW}, Heterochromatin/Repetitive/Copy Number Variation = {Quies, Art}. Table 2: Primers used for ATAC library amplification

### Chromatin immunoprecipitation (ChIP)

U-2-OS cells were transfected in biological duplicate with siRNAs (siCont and siTLK1+2) and 48 hours post-transfection were harvested by trypsinization. Cells were crosslinked with 1% Formaldehyde in DMEM for 10 min at room temperature. Excess formaldehyde was quenched with Glycine at a final concentration of 125 mM. Cells were washed with PBS twice, pelleted by centrifugation at 300 g and dry pellets were frozen at −80°C. For nuclear extract preparation, cells were thawed on ice, resuspended in cold Swelling buffer (25 mM Hepes pH 7.9, 1.5 mM MgCl2, 10 mM KCl, 0.1% NP-40, 1x protease (Roche) and phosphatase inhibitors (Sigma-Aldrich)) for 10 min and passed through a douncer 50 times. Nuclei were pelleted by centrifugation at 3000 g for 5 min at 4°C, and resuspended in 300 ul of 1% SDS in ChIP buffer (10 mM Tris-HCl pH 7.5,150 mM NaCl, 1% Triton X-100, 5 mM EDTA, 0.5 mM DTT, 1x protease (Roche) and phosphatase inhibitors (Sigma-Aldrich)). The extracts were incubated for 15 min on ice, changed to 1.5 ml Bioruptor Pico Microtubes (Diagenode) and sonicated in a Bioruptor Pico sonication device (Diagenode) for 30 cycles 30” on/30” off. Chromatin was cleared by centrifugation at top speed 15 min at 4°C and recovered in a low binding tube. Chromatin was checked for size after a phenol/chlorophorm extraction by an agarose gel and by a Bioanalyzer DNA HS Chip (Agilent; Integrated Sciences) so chromatin fragments have a size of 200 bp on average. For H3.3 (Millipore 09-838) and H3K9me3 (Abcam ab8898) ChIP, 5 ug of total chromatin was diluted 1:10 in ChIP buffer and incubated with 1 ug of antibody on rotation at 4°C overnight. 50 ul of prewashed Dynabeads Protein G (Thermo Fisher Scientific) were added on rotation for 2h at 4°C. Beads were washed with Low Salt buffer (50 mM HEPES pH 7.5, 140 mM NaCl, 1% Triton X-100, 1x protease (Roche) and phosphatase inhibitors (Sigma-Aldrich)), High Salt buffer (50 mM HEPES pH 7.5, 500 mM NaCl, 1% Triton X-100, 1x protease (Roche) and phosphatase inhibitors (Sigma-Aldrich)) and eluted by incubating in a thermomixer with Elution buffer (1% SDS, 100 mM NaHCO3) for 30 min at 65°C and 1000rpm. Samples were reverse-crosslinked by incubating at 65°C overnight and incubated with Proteinase K for 1h at 45°C. Chipped DNA was purified using the MinElute PCR Purification Kit (Qiagen) and eluted in 40 ul. Purified ChIP DNA was used for library generation using the NEBNext Ultra II DNA Library Prep Kit for Illumina (NEB) following manufacturer’s instructions. Each of the libraries was labeled by a specific barcode provided in NEBNext Multiplex Oligos for Illumina (Index Primers Set 1 and 2) (NEB) and amplified 9-13 cycles (depending on initial material amount) by PCR in the presence of SYBR Green in order to obtain an optimal yield. Libraries were quantified using DNA HS Qubit (Thermo Fisher Scientific) and size measured by a 2100 Bioanalyzer Instrument (Agilent) and a DNA HS Chip (Integrated Sciences). An equimolar pool was generated with the eight libraries and the pool was sequenced in 2 lanes 50 nt single read in an Illumina HiSeq2500 (Illumina). Libraries were also used as a template for qPCR using the primers corresponding to telomeric repeats (tel) and HERV summarized in Table 3.

For ChIP-seq analysis, single end reads of 50 bp length were aligned to the hg19 human genome version using Bowtie v0.12.9 (Langmead et al., 2009) with default parameters. Alignments were sorted and indexed using Sambamba v0.5.8 (http://lomereiter.github.io/sambamba/). Duplicated reads were removed using Sambamba. Peaks were called with MACS v1.4.2 (Zhang et al., 2008) for each sample with the corresponding input as control. All parameters were left as default except for read length. Peaks were annotated with the HOMER v2.8.2 (Heinz et al., 2010) software with the hg19 annotations. Consensus peaks were defined as the union of all peaks from all samples. For each sample, the number of reads per peak was computed using the countBam function from the Rsamtools R package (version 1.30.0, http://bioconductor.org/packages/release/bioc/html/Rsamtools.htm). Differential binding was computed using DESeq2 (Love et al., 2014) with replicate as covariate. Normalized densities for peaks were computed using the annotatePeaks function from the HOMER suite. Densities were plotted using the feature AlignedDistribution function from the ChIPpeakAnno R package (Zhu et al., 2010).

### RNA-Seq

U-2-OS cells were transfected in biological duplicate with siRNAs (siCont, siTLK1, siTLK2, siTLK1+2) and 48 hours post-transfection were washed twice with cold PBS and collected by scraping in Tri-Reagent (Sigma-Aldrich). RNA was isolated using the PureLink RNA Mini Kit (Thermo Fisher Scientific) following the manufacturer’s instructions for TRIzol Plus Total Transcriptome Isolation. Briefly, chloroform extraction followed by centrifugation resulted in a colorless upper aqueous phase that was mixed 1:1 with 100% ethanol. Sample containing RNA in 50% ethanol was bound to the spin cartridge, washed twice with Wash Buffer II and eluted in RNase-free water.

To avoid limiting our analysis in mRNA polyA+, we performed enrichment of whole transcriptome RNA by depleting ribosomal RNA (rRNA) species. Purified RNA samples were first quantified for integrity, size and purity by a 2100 Bioanalyzer Instrument (Agilent) and an RNA 6000 Nano Chip (Integrated Sciences). 2 ug of total RNA were processed for rRNA depletion by Ribozero Gold rRNA Removal kit (Human/Mouse/Rat) (Illumina) following manufacturer’s instructions. rRNA-free RNA was subsequently fragmented and cDNA generated by NEBNext RNA First Strand Synthesis Module (NEB) and NEBNext mRNA Second Strand Synthesis Module (NEB). Resulting cDNA was purified, quantified and used for library generation for Next Generation Sequencing using the NEBNext Ultra II DNA library prep for Illumina (NEB). Each of the libraries was labeled by a specific barcode provided in NEBNext Multiplex Oligos for Illumina (Index Primers Set 1) (NEB) and amplified 7 cycles by PCR in the presence of SYBR Green in order to obtain an optimal yield. Libraries were quantified using DNA HS Qubit (Thermo Fisher Scientific) and size measured by a 2100 Bioanalyzer Instrument (Agilent) and a DNA HS Chip (Integrated Sciences). An equimolar pool was generated with the eight libraries and the pool was sequenced in 2 lanes 50 nt single read in an Illumina HiSeq2500 (Illumina).

Single end reads of length 50 bp were aligned to the human genome version hg19 using bowtie v0.12.9 (Langmead et al., 2009) with default parameters. Alignments were sorted and indexed using Sambamba v0.5.8 (http://lomereiter.github.io/sambamba/). Gene differential expression was performed using DESeq2 (Love et al., 2014) with replicates as covariate. Pathway enrichment was assessed through the preranked version of Geneset Enrichment Analysis (GSEA)(Subramanian et al., 2005). GSEA was applied to the ranking defined by the log2 Fold Change of the differential expression analysis using DESeq2. Genesets for analyses were from the Gene Ontology (GO) terms (Ashburner et al., 2000) as collected in the GSEABase R package (version 1.40.1), or from the Hallmark collection (Liberzon et al., 2015) after retrieval from the MsigDB(Liberzon et al., 2011). Classification in coding and non-coding gene classes was performed according to the Gencode annotation version 19(Harrow et al., 2012). For Repeat Masker analysis, sequences from repeat elements in the human Repbase database version 22.06 (Bao et al., 2015) were downloaded. Reads were aligned using bowtie with parameters –best and -k=1. Fold changes were computed between siTLK1+2 and siCont samples using normalised reads values of those repeat sequences that had at least 100 raw reads.

### C-circle assay

The C-circle assay protocol was adapted from(Henson et al., 2017). Genomic DNA from 200 000 cells was extracted by incubating cells with 50 ul of QCP lysis buffer (50 mM KCl, 10 mM Tris-HCl pH 8.5, 2 mM MgCl2, 0.5% IGEPAL CA-630, 0.5% Tween-20) and 3 ul of QIAGEN protease shaking at 1400rpm at 56°C for 1 hour. The QIAGEN protease was inactivated by incubating the samples at 70°C for 20 min. DNA concentration was measured by fluorimetry using the Qubit dsDNA HS Assay (Thermo Fisher Scientific). Samples purified from HeLa LT cells were pre-diluted in QCP lysis buffer at 30 ng/ul. 30 ng of DNA were diluted to 10 ul in 10mM Tris-HCl pH 7.6 and mixed with 9.25 ul of Rolling Circle Master Mix (RCMM) (8.65mM DTT, 2.16X 10X φ29 Buffer, 8.65ug/mL BSA, 0.216% Tween-20 and 2.16mM of each dATP, dCTP, dGTP and dTTP) and 0.75 ul of φ29 DNA Polymerase (NEB). Rolling Circle Amplification was performed by incubating samples in a thermocycler at 30°C for 8 hours, polymerase was inactivated at 70°C for 20 min and then kept at 8-10°C. Samples were kept at −20°C. For slot blot detection, samples were diluted with 2x SSC to 200ul, then slot-blotted onto Nytran SuPerCharge (SPC) nylon blotting membranes (Sigma-Aldrich) under native conditions. After 254 nm UV-C crosslinking, the membrane was hybridized with γ-32P labeled Tel-C oligo probe (CCCTAA)4 in hybridization buffer (1.5X SSPE, 10% polyethylene glycol (PEG) MW 8000, 7% SDS) for 16h. Membrane was exposed onto a storage phosphor screen (Molecular Dynamics) and scanned using Typhoon 8600 Variable Mode Imager (Molecular Dynamics). Membrane was stripped in wash solution (0.5X SSC, 0.1% SDS) at 65 °C and re-hybridized with γ-32P labeled Alu oligo probe 5’-GTAATCCCAGCACTTTGG-3’ for loading control.

### Analysis of TCGA signatures

TCGA RNA-Seq datasets were downloaded from the legacy archive of the NCI GDC commons database (Grossman et al., 2016) and processed separately for each cancer type. Expression measures were expressed in RSEM (Li and Dewey, 2011) in this TCGA version, which were log2-transformed and quantile normalized. For patients with multiple instances, a single sample chosen at random was kept while the rest were excluded from further analyses. For a number of cancer types there were two different platforms available: Illumina HiSeq 2000 (HiSeq) and Illumina Genome Analyzer (GA); in such cases, duplicated samples across platforms were removed from the GA dataset while the HiSeq instance was kept in the final dataset. Samples whose expression values showed an unusual distribution compared to the rest of samples in their datasets were also excluded (ACC: TCGA-OR-A5L9-01A; HNSC: TCGA-D6-A6ES-01A and TCGA-CV-A45Q-01A; LAML: TCGA-AB-2955-03A, TCGA-AB-2986-03A, TCGA-AB-2816-03A, TCGA-AB-2955-03A, TCGA-AB-2986-03A and TCGA-AB-2816-03A; LIHC: TCGA-DD-A3A6-11A and TCGA-FV-A4ZP-01A; SKCM: TCGA-D3-A2JK-06A; UCEC: TCGA-BS-A0V4-01A; UVM: TCGA-WC-A885-01A; COAD: TCGA-A6-2679-01A and TCGA-AA-A004-01A). For each cancer type, expression matrices were corrected a-priori by platform, source center and plate id, when suitable. For doing so, a linear model was fitted to the expression values gene wise in which platform and source center were included as fixed effects. Regarding sample’s plate id, it was included in the models as a fixed or a random effect depending on the number of levels and sample size available for the cancer type under consideration. In some cases, additional variables were also included in the models in order to preserve signal with biological or clinical relevance (BLCA: diagnosis subtype; BRCA: PAM50 subype; COAD and STAD: microinstability status; KIRP: tumor type).

The T cell CD8, NK cells activated, Macrophages M1 scores, were obtained from (Thorsson et al., 2018) and matched to expression through the sample id. Aneuploidy score and Stromal and Leukocyte fraction estimates were obtained from(Taylor et al., 2018). To estimate chromosomal instability (CIN) we computed signatures with the geneset CIN25 published in(Carter et al., 2006). The interferon signature was taken from our RNA-seq analysis. The genes in the TNFalpha signature were taken from the Hallmark gene set “TNFA_SIGNALING_VIA_NFKB” (Liberzon et al., 2015). ALT status was taken from(Marzec et al., 2015).

Gene signature scores were computed as the mean of all genes in the signature after scaling the expression matrix gene wise. The global signature was defined as the score associated with the gene signature containing all genes in the expression matrix. Correlation coefficients and p-values were computed using the ‘pcor.test’ function from the ppcor R package(Kim, 2015). In order to avoid spurious correlations due to technical and global effects(Caballé Mestres et al., 2018), partial correlations were computed whenever a gene signature was involved with the global signature as adjusting variable.

For the association between ALT status and signatures, expression matrices for all available TCGA datasets were scaled gene wise and merged. Only those cancer types with more than two ALT+ patients were included in the merged dataset. In order to account for possible technical and global effects, gene signature scores were adjusted by the global signature before plotting(Caballé Mestres et al., 2018). P-values and coefficients of the association between gene expression or gene signatures and ALT status were computed through a linear model with the dataset as covariable. Whenever gene signatures were being tested, the global signature was also included as covariable. The scores associated to T cells CD8, stromal fraction, leukocyte fraction and aneuploidy were transformed using the square root to ensure normality of the data for the linear model. P-values were adjusted for multiple comparisons using Benjamini-Hochberg.

### Data availability

The authors declare that the computer code generated during the current study is available from the corresponding author on reasonable request. ATAC-seq data from U-2-OS and HeLa LT, H3.3 ChIP-seq data from U-2-OS and RNA-seq data from U-2-OS have been submitted to the Genbank GEO database (http://www.ncbi.nlm.nih.gov/geo/) and we are waiting for the accession codes. All other remaining data are available within the Article and Supplementary Files, or available from the authors upon request.

### Quantification and statistical analysis

Statistical significance of non-sequencing experiments was determined with the tests stated in the figure legends using PRISM software (Graphpad Software Inc.). All data are from at least two independent experiments. Specific replicate numbers (N) for each experiment can be found in the corresponding figure legends. Statistically significant differences are labeled with one, two, three or four asterisks if p < 0.05, p < 0.01, p < 0.001 or p < 0.0001, respectively. Statistical analysis of TCGA data sets is described in detail in the corresponding methods section.

## Acknowledgements

The authors wish to thank members of the Stracker lab and S.J. Boulton for support and input, N. Pons, D. Fernandez and N. Plana from the IRB Functional Genomics core for library preparation for RNA-seq and ChIP-seq experiments, A. Lladó, L. Bardia and S. Tosi from the IRB Advanced Digital Microscopy facility for ImageJ macro writing, the A. Jordan lab for primers, TBK1i and advice to address repeat/IFN transcription, E. Lazzerini-Denchi for C-circle assay protocols, and the S. Peiro lab for advice on ATAC-seq. HeLa LT and GM847 cells were a kind gift from J. Jacobs/J. Karlseder and J. Lingner, respectively. THS was funded by the Spanish Ministry of Science, Innovation and Universities (BFU2015-68354/GENPATH, GINDATA and FEDER), the Centres of Excellence Severo Ochoa award and the CERCA Programme. SSB was funded by a PhD fellowship and the project LCF/PR/GN14/10270002 from the “la Caixa” Foundation.

## Author contributions

SSB and THS designed the study, interpreted the data and cowrote the manuscript. SSB performed experiments and analyzed data. MVP contributed to reagent generation. CSOA analyzed sequencing and tumor data. THS and SSB secured funding, and THS supervised the project.

## Supplementary Figure Legends

**Figure S1.**
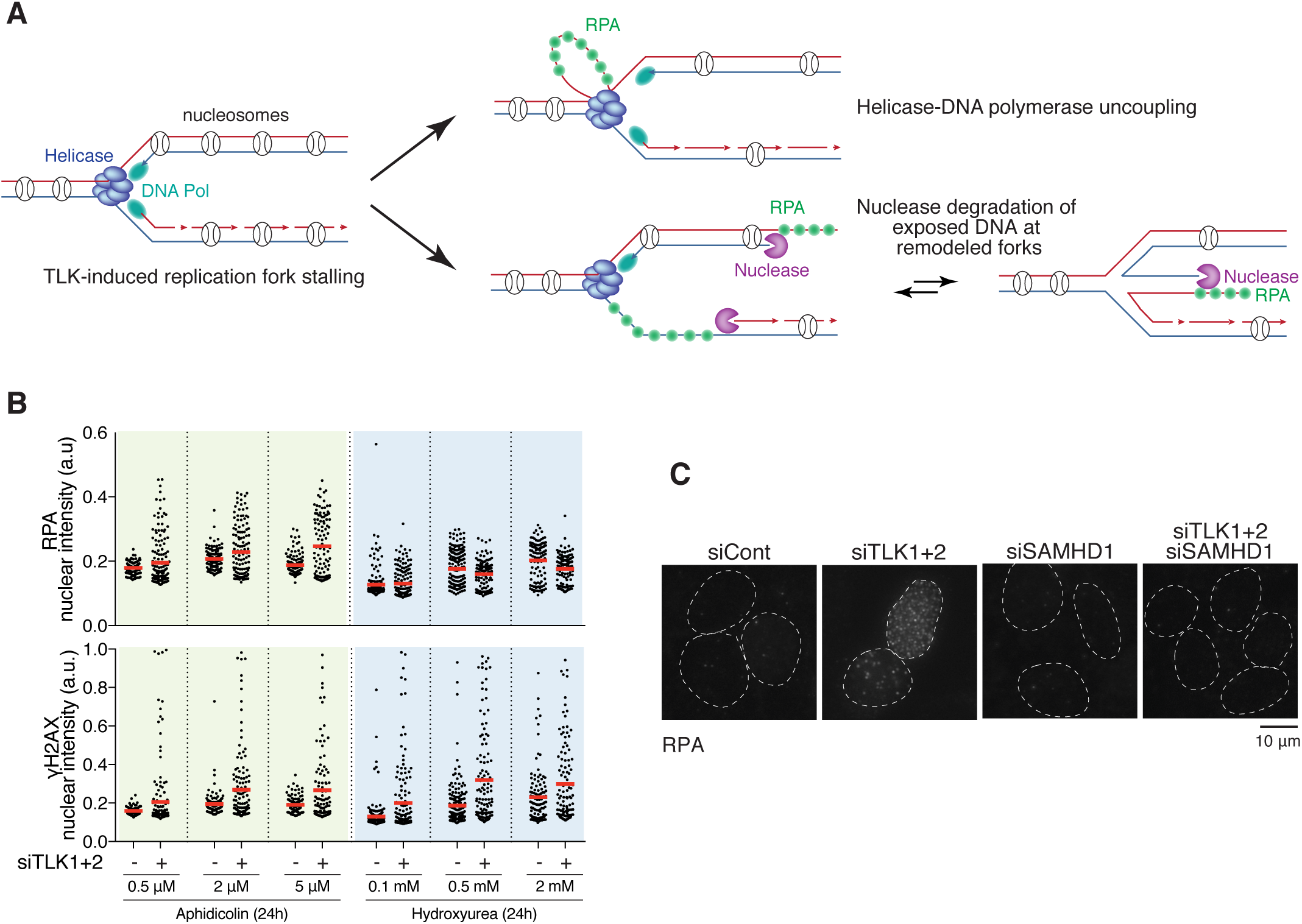
**(A)** Schematic of the question on whether ssDNA accumulation may occur ahead or behind the replication fork in TLK depleted cells. **(B)** HTM quantification of the nuclear intensity of chromatin-bound RPA (top panel) and γH2AX (bottom panel) in U-2-OS cells 48 hours after siRNA transfection and treated with the dose/time indicated. Red bars indicate the average. **(C)** Representative IF of chromatin-bound RPA staining in U-2-OS cells 48 hours after siRNA transfection.

**Figure S2.**
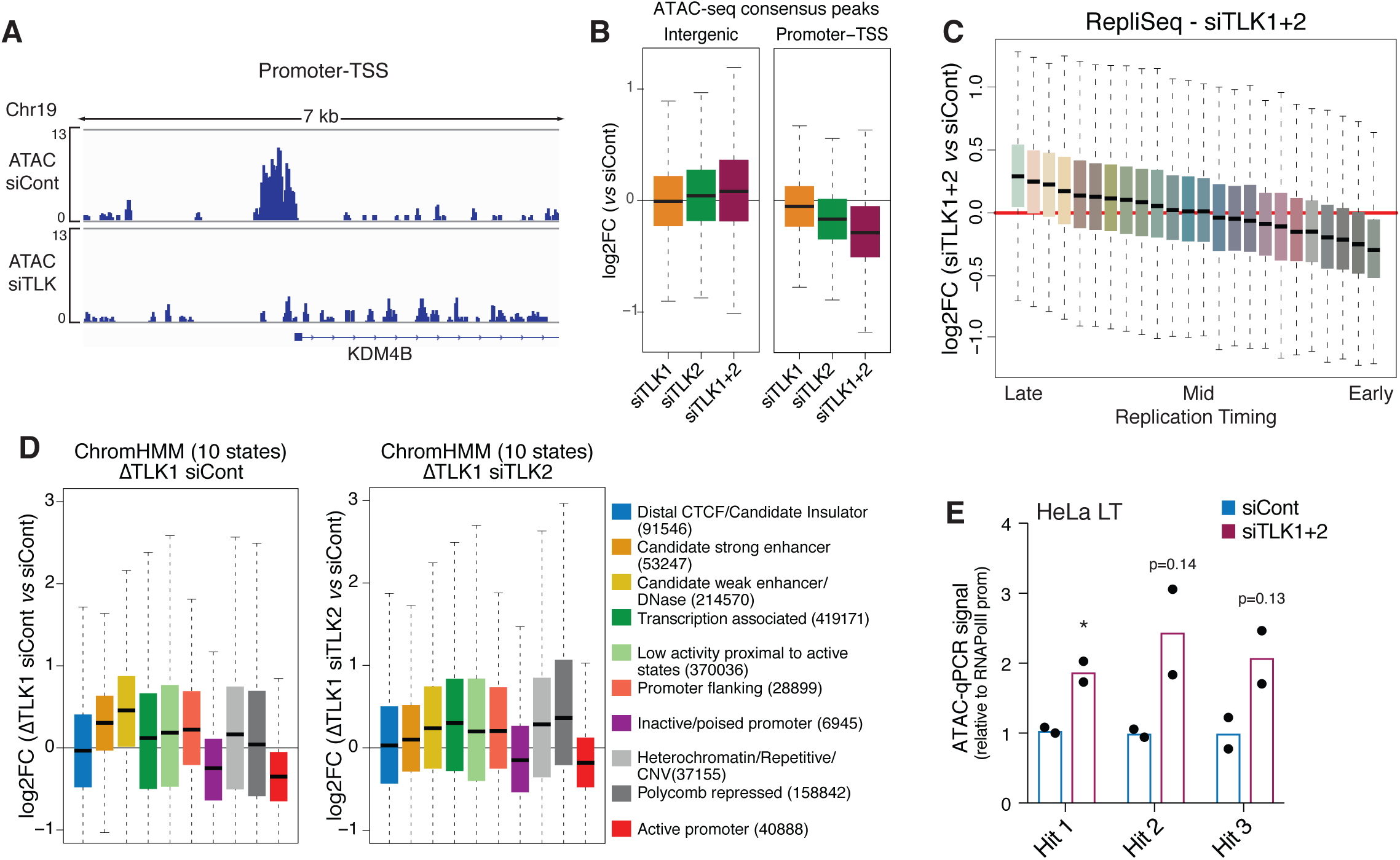
**(A)** Representative IGV track of ATAC-seq reads of a promoter-TSS region. **(B)** Boxplots of ATAC-seq FC (siTLK1, siTLK2, siTLK1+2 relative to siCont) computed in different genomic annotations, namely intergenic and promoter-TSS regions. **(C)** Boxplots of ATAC-seq FC (siTLK1+2 relative to siCont) through different replication timing chromatin regions from NHEK RepliSeq ENCODE dataset (NHEK was used an average track). **(D)** Boxplots of ATAC-seq FC in HeLa LT cells (ΔTLK1 siCont and (ΔTLK1 siTLK2 relative to siCont) through different ChromHMM chromatin states. **(E)** ATAC-qPCR at selected genomic regions in HeLa LT cells as in Figure 2G (n=2).

**Figure S3.**
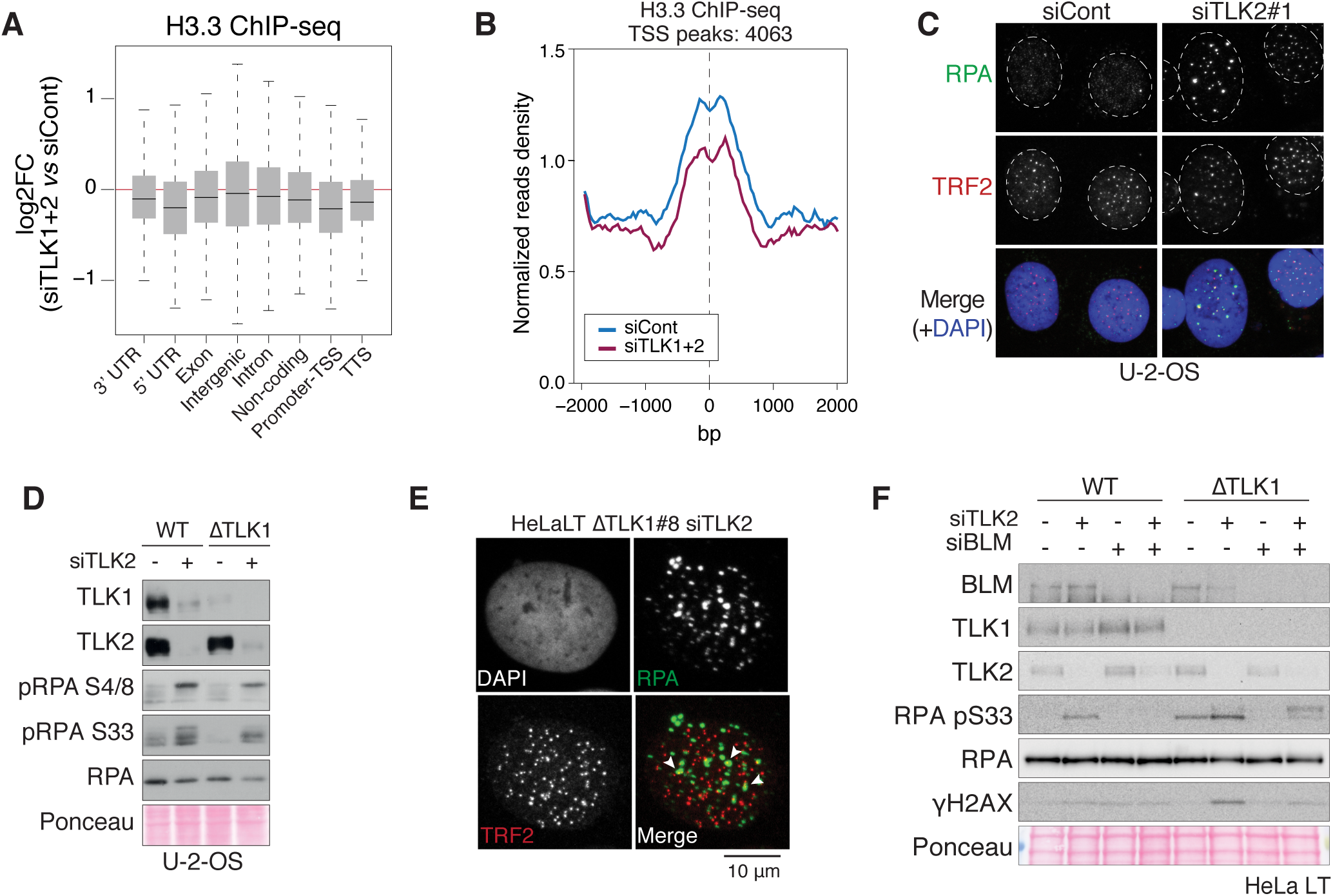
**(A)** Boxplots of H3.3 ChIP-seq FC (siTLK1+2 relative to siCont) through different genomic annotations in U-2-OS cells (n=2). **(B)** Normalized read density at H3.3 peaks located at TSS with +/− 2 kb. **(C)** Representative IF of RPA-TRF2 staining in U-2-OS cells. **(D)** Western blot of replication stress signalling markers upon TLK loss in U-2-OS cells, parental (WT) and TLK1 CRISPR knockout clone (ΔTLK1). **(E)** Representative IF of RPA-TRF2 staining in HeLa LT ΔTLK1 siTLK2 cells. White arrowheads indicate colocalization of RPA-TRF2 foci. **(F)** Western blot of HeLa LT parental (WT) and TLK1 CRISPR knockout clones (ΔTLK1) 48 h after being treated with the corresponding siRNAs.

**Figure S4.**
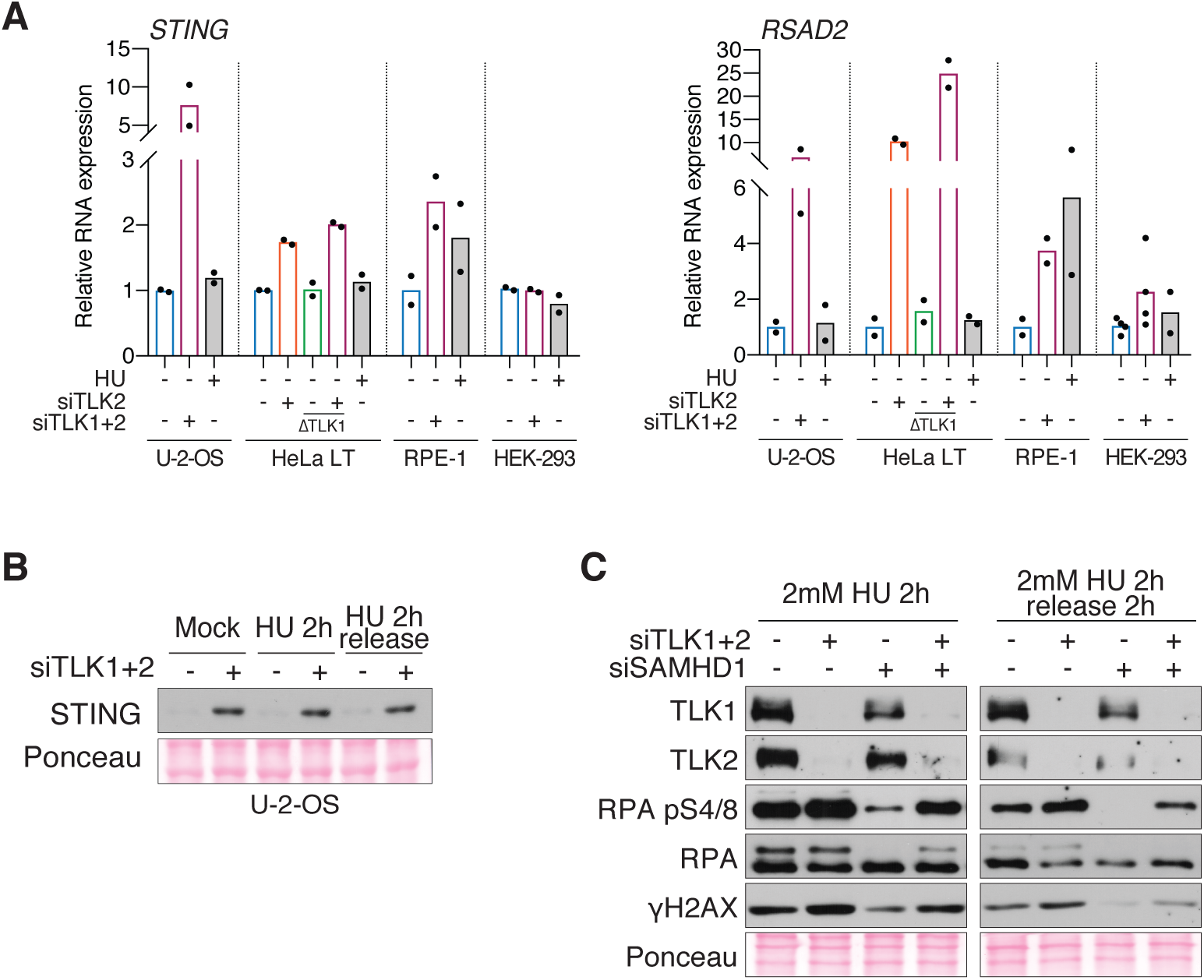
**(A)** Expression levels of STING and RSAD2 by RT-qPCR in U-2-OS, HeLa LT, RPE-1 and HEK-293 cells 48 h after being treated with the corresponding siRNAs, mock treated or treated with 2 mM HU for 2 h. Data were normalized to unchanging expression gene levels (B-actin) and the signal obtained in siCont conditions was set to 1 (n≥2). **(B)** Western blotting of STING in U-2-OS cells, 48 h after being treated with the corresponding siRNAs and HU as in Figure 1B. To evaluate the status of other markers, refer to Figure 1B. **(C)** Western blot showing single or double depletion of TLKs and SAMHD1 in U-2-OS cells treated with 2 mM HU for 2 hours and harvested without a release or with a release in fresh media for 2 hours.

